# Prostate-specific membrane receptor PPAP facilitates *E. coli* invasion of luminal prostate cells via FimH binding

**DOI:** 10.1101/2024.10.28.620623

**Authors:** Maria Guedes, Amruta Joshi, Sina Dorn, Simon Peters, Tristan Beste, Alexander M Leipold, Mathias Rosenfeldt, Antoine-Emmanuel Saliba, Ulrich Dobrindt, Charis Kalogirou, Carmen Aguilar

**Affiliations:** Host Pathways in Urinary Tract Infections Group, Institute of Molecular Infection Biology, University of Würzburg, Würzburg, Germany; Helmholtz Institute for RNA-based Infection Research, Helmholtz-Center for Infection Research, Würzburg, Germany; Institute of Molecular Infection Biology, Faculty of Medicine, University of Würzburg, Würzburg, Germany; Institute of Pathology, University of Würzburg, Würzburg, Germany; Institute of Hygiene, University of Münster, Münster, Germany; Department of Urology and Pediatric Urology, University Hospital Würzburg, Würzburg, Germany

## Abstract

Bacterial prostatitis caused by uropathogenic *Escherichia coli* (UPEC) strains is a highly prevalent and recurrent infection responsible for significant morbidity in men. However, the molecular pathogenesis of prostatitis remains poorly understood, partly due to the lack of comprehensive *in-vitro* models. In this study, we introduce a murine prostate organoid model that replicates the cellular heterogeneity of the prostate epithelium with a cell composition and transcriptional signature comparable to the native prostate tissue. Using this model, we uncovered that UPEC preferentially attaches to, invades, and replicates within luminal prostate cells. This selective interaction is mediated by the binding of the bacterial adhesin FimH to the prostate-specific membrane protein PAPP, which is exclusively expressed on luminal prostate cells. Altogether, we identified a new mechanism by which UPEC infects the prostate epithelium, highlighting FimH’s adaptability in engaging host receptors and its potential for targeted therapeutic strategies.

## INTRODUCTION

Urinary tract infections (UTIs) are among the most common bacterial infections worldwide, with a global incidence of over 400 million cases per year^1^. Despite adequate antibiotic treatments, these infections are highly recurrent, with more than 50% of patients experiencing a recurrence within a year, indicating that current treatment options are less than ideal. While women encounter a higher incidence of UTIs, infections in men are more challenging to treat, often requiring prolonged and higher concentrations of antibiotics^2,3^. One of the most common complications in male UTIs is the spread of the infection to the prostate gland^4^. Bacterial prostatitis affects nearly 1% of men worldwide, with a bimodal distribution among younger and older men^5^. Due to the proximity of the prostate to the bladder, a shared infection route, common causative infective agents, as well as risk factors, bacterial prostatitis is increasingly considered to be a UTI^6^. As in the bladder, uropathogenic *Escherichia coli* (UPEC) is the most common cause of bacterial prostatitis (up to 80% of all cases)^6–8^. However, very little is known regarding UPEC’s pathogenesis in prostate tissue. In the bladder, once the bacteria reach the lumen, UPEC uses type 1 pili with the FimH adhesin at their tips to bind to the membrane protein uroplakin 1a (UPK1a) of the superficial urothelial cells. After entering these cells, UPEC starts to replicate and subsequently forms intracellular bacterial communities (IBCs)^9,10^. Superficial cells containing IBCs are quickly exfoliated, exposing the underlying immature transitional epithelial cells^9^. FimH can also bind to α3β1 integrins on those cells and aid its invasion to form quiescent intracellular reservoirs (QIRs)^11,12^, which are responsible for antibiotic tolerance and recurrences. Understanding whether these characteristics of infection in the bladder are also seen in the prostate tissue, is fundamental, as it influences the design and implementation of treatments.

The prostate is a male sex gland that contributes roughly 30% of the seminal fluid. It is made up of ducts lined with pseudostratified epithelium composed of three main types of cells: (1) secretory luminal cells expressing keratin (KRT) KTR18 and androgen receptor (AR); (2) basal cells, characterised by their expression of KRT5, KRT14, and p63; and (3) rare neuroendocrine cells, positive for chromogranin A (CHGA)^13^.

Despite the clinical relevance of bacterial prostatitis, prostate infections have been significantly understudied. This is partly due to technical difficulties in infecting *in vivo* mouse models and the lack of suitable *in-vitro* infection models mimicking the cell heterogeneity of the prostatic epithelium. A few studies have examined prostate infections using mouse models, and, to our knowledge, none have utilised primary cells from human or mouse tissues to investigate these infections. Collectively, mice studies have shown that UPEC, either prostate isolates (e.g., CP1) or cystitis isolates (e.g., UTI89) of UPEC, can establish acute and chronic infection in the murine prostate gland, as well as induce a robust proinflammatory response, immune cell infiltration, and hyperplasia in the tissue^14–21^. Interestingly, one of those studies observed a UPEC tropism for the prostate gland versus the bladder in C57BL/6 male mice^20^. The same study showed that similar to the male bladder, the murine prostate fails to induce a protective immune response to the infection, leading to chronic prostatitis. However, how the bacteria interact with epithelial cells and whether or not they invade and/or replicate within these host cells remains unknown.

Over the past decade, organoids and organoid-based models have emerged as powerful *in-vitro* systems, replicating key physiological and functional characteristics of their corresponding organs. Derived from tissue-resident adult stem cells, these models can differentiate into the various epithelial cell types found within the tissue, closely mimicking the structure, function, and cellular complexity of the organ. Prostate organoids, in particular, have proven valuable for studying prostate development and cancer^22,23^. However, a suitable model for investigating host-pathogen interactions is still lacking.

Here, we harness the power of organoids to develop a model of the prostate epithelium to study UPEC pathogenesis in the context of bacterial prostatitis. Our model closely mimics the cellular heterogeneity of the epithelial compartment in the real organ. Using this model, we found that UPEC adheres to, invades, and replicates within luminal prostate cells. This interaction was dependent on the binding of the FimH adhesin to the newly identified prostate-specific membrane receptor Prostatic Acid Phosphatase (PPAP).

## RESULTS

### An organoid-based model of the murine prostate closely mimics the tissue’s epithelial heterogeneity

In general, a suitable *in-vitro* infection model should reflect the cellular complexity of the tissue it is developed to represent. In the case of a prostate infection model, this should include the main cell types (i.e. luminal gland cells and basal stem cells) present in the prostate epithelium with the exception of the rare neuroendocrine cells. To set up such an *in-vitro* model, we started by generating mouse prostate 3D organoids from C57BL/6 wild-type mice following a previously published protocol^24^. However, to use the model for UPEC infections, the apical surface of the epithelium should be easily accessible, which is not the case for 3D organoids grown in extracellular matrix. Thus, we seeded the organoid cells into a 2D surface (**Figure 1a**) using the standard prostate organoid media previously described^24^. However, these conditions did not achieve a very high degree of differentiation, as observed by the low number of differentiated luminal cells positive for Cd24a and the high number of Krt5*-*positive basal cells (**Figure 1b**, middle panel). Given that activation of the androgen receptor pathway is the main driver for prostate epithelium development and differentiation, we hypothesised that increasing the androgen levels in the medium could improve cellular differentiation. Increasing the concentration of 5α- dihydrotestosterone (DHT) from 1nM to 10nM was sufficient to increase the differentiation of basal cells into luminal cells (**Figure 1b**, right panel). In contrast, removing DHT from the medium enriched the model for basal stem cells (Krt5-positive cells; control condition; **Figure 1b** left panel).

**Figure 1.**
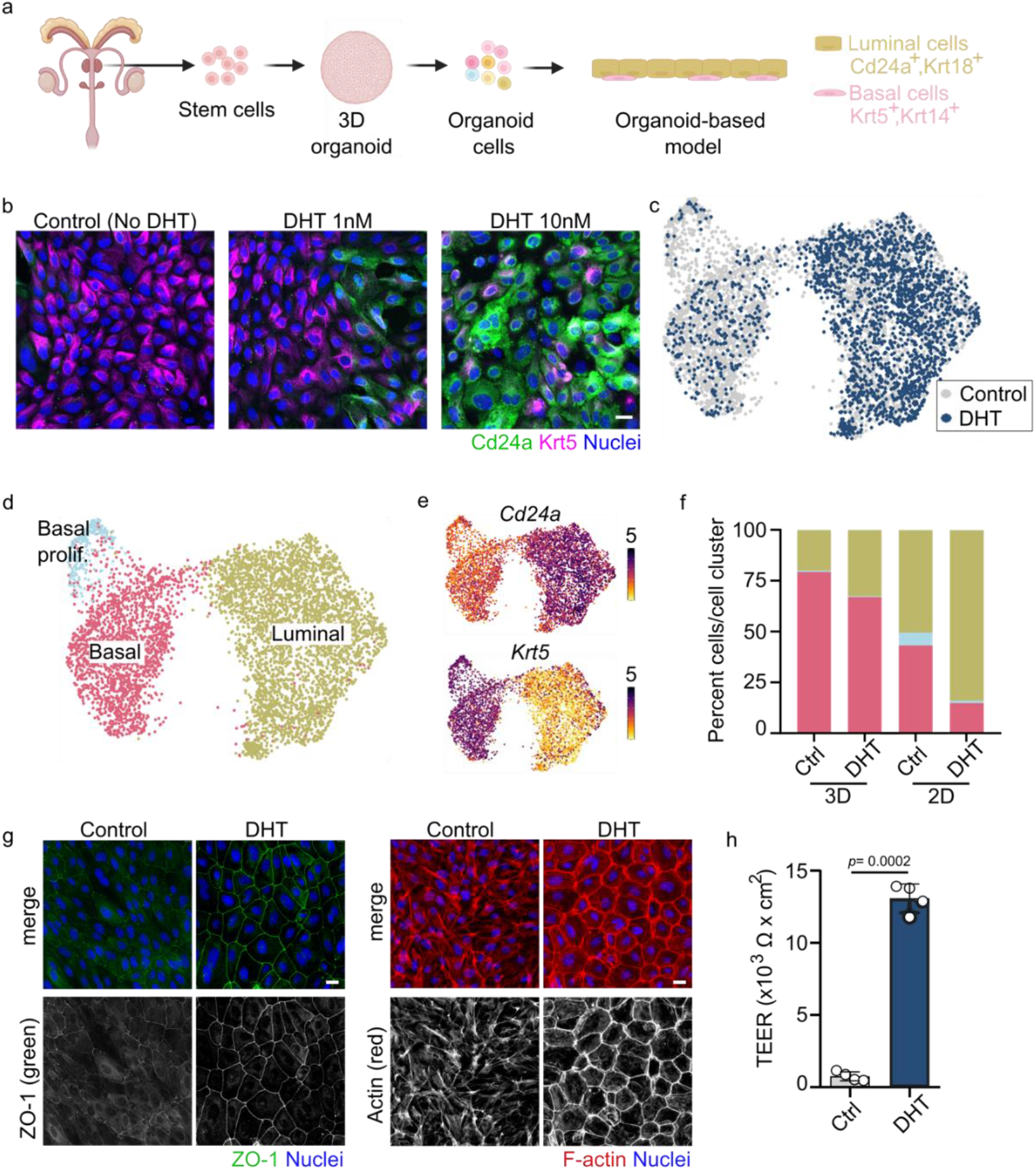
An organoid-based model of the murine prostate reflects epithelial cell heterogeneity. **a.** Scheme of the experimental setup. Prostate organoids were generated from murine prostate tissue and seeded in 2D. **b.** Confocal microscopy images of the organoid-based model grown in the presence or absence of DHT (1 or 10 nM) and stained for Cd24a (green) and Krt5 (magenta) using immunofluorescence (*n* = 3). Nuclei were counterstained using Hoechst 33342 (blue). Scale bar: 25 µm. **c.** Single-cell transcriptomes from the 2D organoid-based models grown in control (3,602 cells; grey) or DHT (10 nM, dark blue) medium (2,127 cells) were integrated and projected using Uniform Manifold Approximation and Projection (UMAP). **d.** Using known prostate marker genes for the specific prostate cell types, cells have been clustered on the UMAP and assigned to their identities. **e.** Expression of known markers specific for luminal prostate cells (*Cd24a*) and basal cells (*Krt5*) colour-coded and projected on top of the UMAP displayed in panels (**c**) and (**d**). **f.** Cell percentage per cluster identified in panel (**d**) and colour coded as in panel (**d**; basal proliferative cells shown in light blue, basal in pink, and luminal in khaki). **g.** Confocal microscopy image of the organoid-based model grown without or with 10 nM DHT and stained for ZO-1 (green; left panel) and F-actin (red; right panel; *n* = 5). Nuclei were counterstained using Hoechst 33342 (blue). Scale bar: 25 µm. **h.** TEER analysis of the 2D organoid-based model grown in the presence or absence of DHT (10 nM; *n* = 4)

To obtain a more in-depth characterisation of our 2D model, we applied single-cell RNA sequencing (scRNA-seq, see Methods section) to characterise the cellular composition in 3D organoids versus the 2D model grown in both culture conditions (control and 10nM DHT). Altogether, we analysed 9992 single-cell transcriptomes (**Supplementary Table 1**). Unsupervised clustering identified three clusters in both the 3D organoid and the 2D model, corresponding to basal cells (positive for *Krt5*, *Trp63*, *Krt14*), luminal cells (positive for *Cd24a*, *Krt8*, *Krt18*, *Psca*) and highly proliferative basal cells (positive for *Mki67* and *Birc5*; **Figure 1c-e**, **Extended Data** Figure 1, **Supplementary Table 2**). However, we observed a higher percentage of luminal cells in the 2D model compared to 3D organoids, independent of the absence/presence of DHT, indicating an overall higher degree of differentiation in the 2D versus the 3D model (20.1% versus 50.5% in Ctrl medium and 83.9% versus 32.6% in DHT medium, respectively; **Figure 1f**, **Supplementary Table 1**). Moreover, supplementation with 10nM DHT increased the number of luminal cells in both models, with a more substantial effect in 2D (from 50.5 to 83.9%) than in 3D (20.1% to 32.6%; **Figure 1f**). In addition, a neighbourhood analysis of the 3D and 2D model data using the murine prostate tissue data from previous studies^25,26^ as reference showed that the 2D model grown with DHT had the highest transcriptional similarity at the single cells level with the primary tissue (**Extended Data** Figure 2). Of note, the number of luminal cells in the 2D model grown in the DHT medium (83.9%) was also reminiscent of the numbers observed in murine tissue (**Supplementary Table 1**)^25^.

The scRNA-seq data also showed that in the 2D model, the presence of DHT leads to increased transcription of cell polarisation and epithelial barrier integrity markers (*Ocln*, *Ctnnb1*, *Cdh1*, *Epcam*, *Tjp1*, *Tjp2*, *Tjp3*, *Cldn1*, *Cldn3*, *Cldn4*, *Cldn7* and *Cldn23*; **Extended Data** Figure 3a-c). However, those markers did not show a clear increase in cell polarisation in the 3D organoids (**Extended Data** Figure 4a-c). Cell polarisation upon differentiation with DHT in the 2D model was confirmed by immunostaining of the Zonula occludens 1 (ZO-1) protein and the actin filaments at the cell junctions (apical ring of actin filaments; **Figure 1g**). In addition, an increased transepithelial electrical resistance (TEER) was measured in the differentiated 2D model (**Figure 1h**), indicating an overall increase in epithelial barrier function. Together, our data demonstrate that the differentiation of prostate organoid cells can be directed *in vitro* to mimic the epithelial compartment of the murine prostate epithelium.

### UPEC preferentially invades luminal prostate cells

Invasion and replication within host cells are crucial steps for many pathogenic bacteria since they allow them to avoid clearance by the immune response and/or antibiotic treatment. For UPEC, different infection/survival strategies have been described: in the bladder, UPEC can invade and replicate very efficiently within the differentiated superficial umbrella cells^10^, whereas in the kidney, it replicates extracellularly to form dense intratubular biofilm-like communities^27^. In the prostate, however, the fate of UPEC remains unknown thus far. Therefore, we first aimed to characterise UPEC’s ability to infect prostate cells using our 2D organoid-based model. Given that most of the *E. coli* pathotypes isolated from bacterial prostatitis are UPEC strains, we used one of the most common reference UPEC strains, UTI89, isolated from a cystitis case^28^.

To distinguish between extra- and intracellular bacteria in the 2D model grown either in the presence or absence of DHT, we constructed a GFP-expressing UTI89 strain and combined with a differential immunostaining protocol where only extracellular bacteria are labelled with an anti-LPS antibody. This differential staining showed that UPEC rapidly invaded the 2D model, with a clear tropism to the cells in the differentiated model (**Figure 2a**). Of note, only live bacteria were able to invade cells, as PFA-killed, heat-killed bacteria or fluorescent beads were not observed intracellularly (**Extended Data** Figure 5a-b).

**Figure 2.**
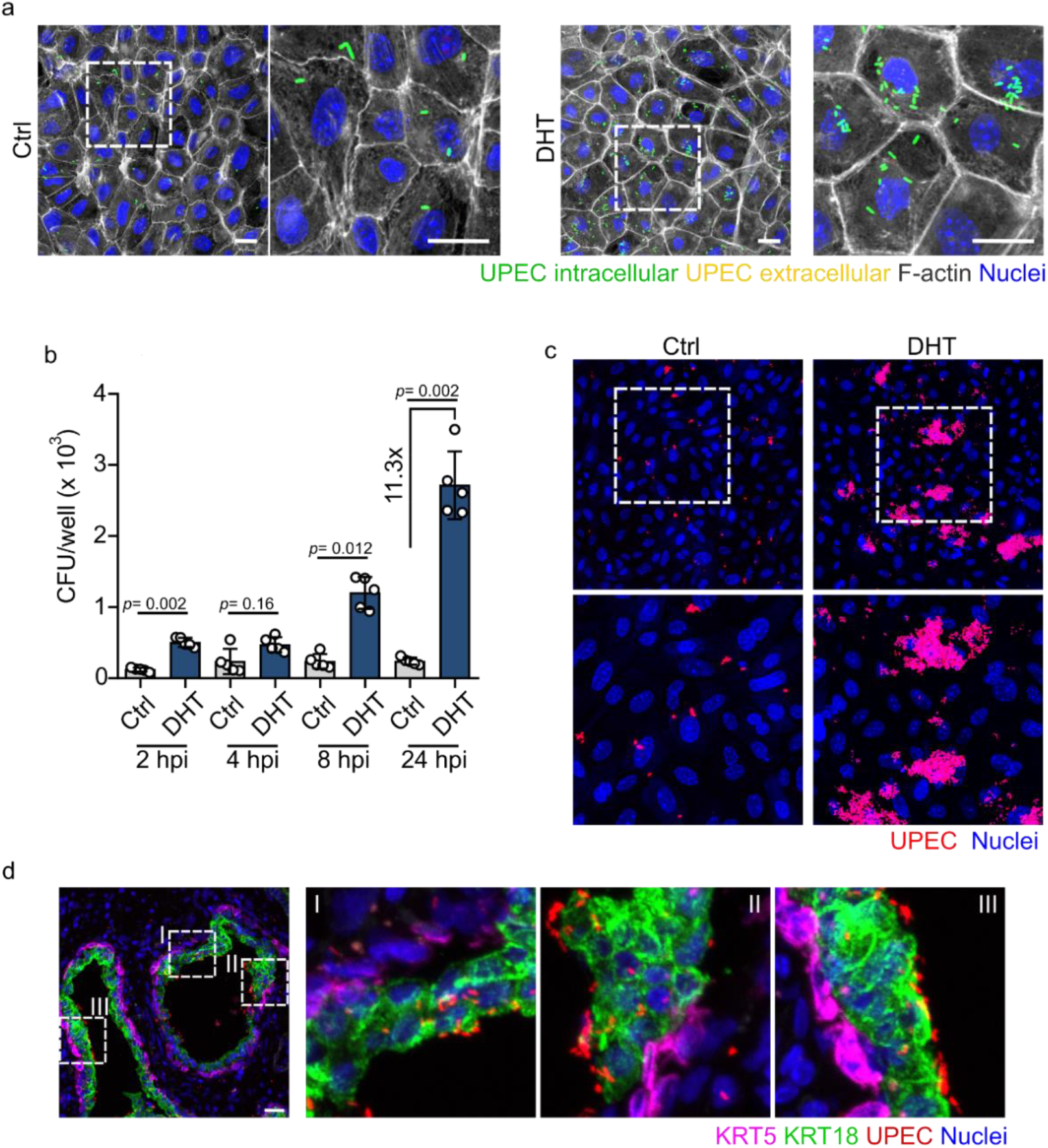
UPEC preferentially invades and replicates in cells within the differentiated 2D organoid-based model. **a.** Confocal microscopy images of the organoid-based prostate model at 1 hour post-infection (hpi) with UPEC strain UTI89 (green). The model was grown without (left, Ctrl) and with 10 nM DHT (right) and stained for F-actin (grey), extracellular bacteria were stained for LPS (red), and nuclei are counterstained with Hoechst 33342 (blue). To differentiate extracellular from intracellular bacteria, cells were stained without permeabilization, which would result in immunostaining of only extracellular bacteria. Extracellular bacteria are shown in yellow (green + red), intracellular in green. Scale bar: 25 µm (*n* = 3). **b.** UPEC colony-forming units (CFUs) were quantified at 2, 4, 8, and 24 hpi in the organoid-based model grown with and without DHT (*n* = 5) **c.** Representative confocal images of the organoid-based model 24 hpi. UPEC is shown in red. Scale bar: 25 µm. **d.** Confocal images of human prostate tissue incubated with UPEC and stained for KRT5 (magenta) and KRT18 (green), nuclei was counterstained using Hoechst 33342 (blue). Scale bar: 25 µm. Data are representative results from 2 donors and 2 independent experiments.

Since the bacteria readily invaded prostatic cells, we next addressed their intracellular fate. Interestingly, we observed that UPEC replicates more efficiently in the differentiated model (DHT; 11.3 fold, **Figure 2b**), enriched for luminal cells, than in the undifferentiated model, enriched for basal prostate cells. In particular, at later time points (24hrs post-infection) it becomes evident that the increased bacterial load in the differentiated model is less likely to stem from an increased initial invasion but rather an increased rate of replication inside the invaded luminal cells (**Figure 2c**). Of note, DHT had no effect on bacterial growth itself (**Extended Data** Figure 6). To validate UPEC’s preference for luminal prostate cells in human tissue, human prostate tissue slides were incubated with UPEC as well. Given that a biopsy is not a common procedure in bacterial prostatitis or male UTI patients, we utilized prostate tissue blocks from patients who had undergone biopsies for other reasons, such as cancer screening. Only tissues identified as healthy by the pathologist were included in the analysis. Microscopy analysis showed a higher number of UPEC attached to the luminal prostate cells (positive for KRT18) than to the basal cells (positive for KRT5; **Figure 2d**). Overall, the results suggest that UPEC invades and replicates effectively in a murine differentiated prostate model enriched with luminal cells and that bacterial binding is also more efficient in human luminal prostate cells.

### FimH is necessary for UPEC invasion into luminal cells

UPEC is a highly versatile pathogen that utilises a variety of adhesins to efficiently infect different environments and/or tissues. Understanding how the bacteria interact with host cells could direct research into anti-adhesive therapies. For example, while type 1 pili are crucial for bladder colonisation, P pili are more significant for adhesion in the kidneys^29,30^. Since the interaction of UPEC with prostatic cells has thus far not been characterised, we aimed to identify UPEC adhesin(s) responsible for bacterial adhesion and invasion in the luminal prostate cells. Given their essentiality in bladder colonisation and the close proximity between the bladder and prostate, we initially investigated the role of type 1 pili, specifically the tip adhesin FimH, in this interaction. Infection of both conditions (Control and DHT) with a FimH deletion mutant showed a clear invasion impairment compared to wild-type bacteria (Δ*fimH*; **Figure 3a-b**).

**Figure 3.**
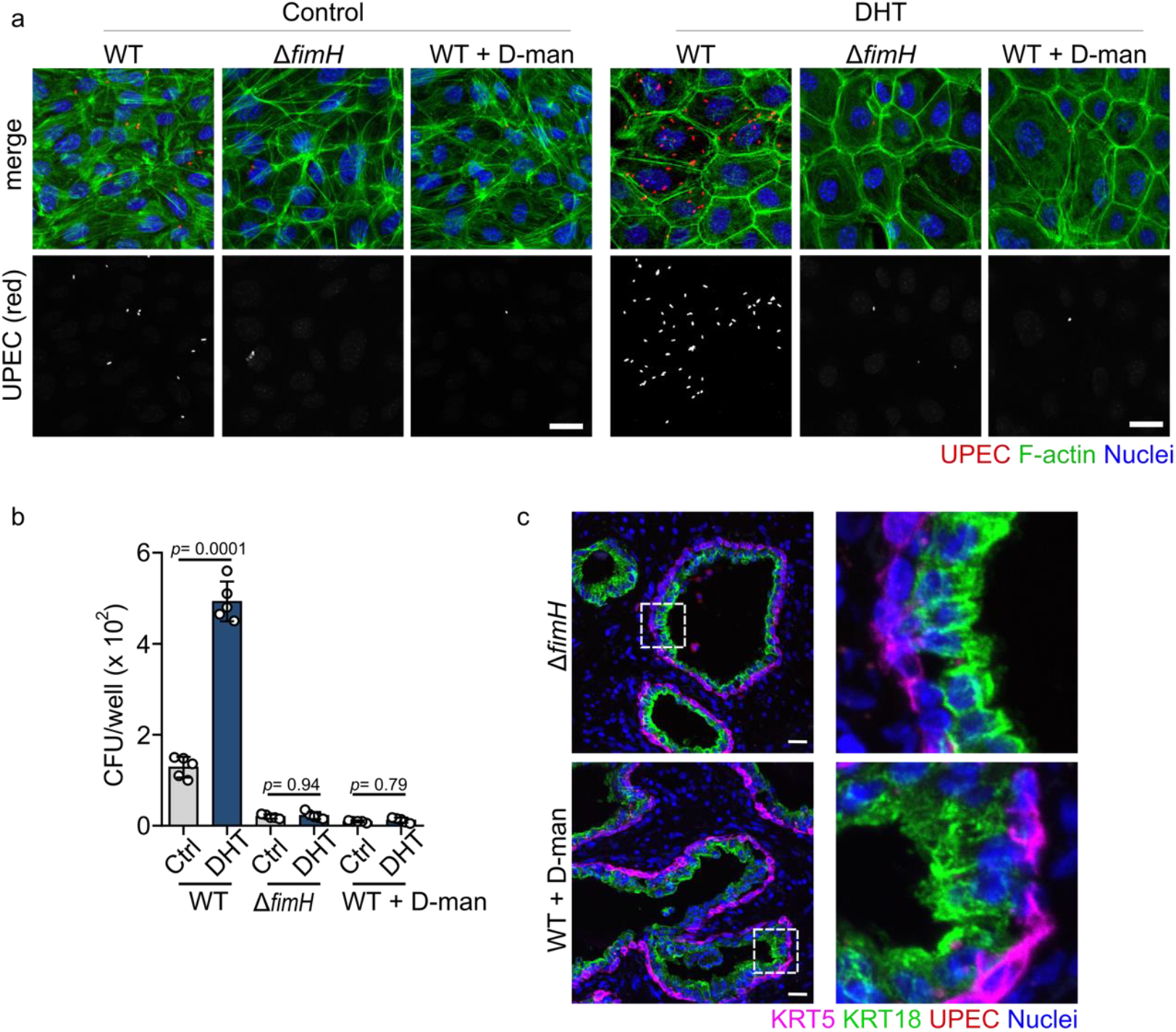
FimH is necessary for maximal invasion into differentiated prostate cells. a-b. Representative images (**a**) and CFUs (**b**) of the organoid-based prostate models infected with WT UTI89, UTI89 Δ*fimH* mutant, and WT UTI89 in the presence of 2.5% D-mannose for 1h. Bacteria are shown in red. Cells were stained for F-actin (green), and nuclei were counterstained with Hoechst 33342 (blue). Scale bar: 25 µm (*n* = 3 for **a**; *n* = 5 for **b**). **c.** Representative confocal images showing binding of Δ*fimH* mutant and WT treated with 2.5% D-mannose (red) to human prostate tissue. The tissue was stained for KRT18 (green) and KRT5 (magenta), and nuclei were counterstained using Hoechst 33342 (blue). Scale bar: 25 µm. Data are representative results from 2 donors and 2 independent experiments.

Moreover, allosteric blockage of the mannose-binding pocket of FimH by adding D-mannose to the infection media resulted in a similar invasion defect of the WT reminiscent of the *fimH* deletion mutant (**Figure 3a-b**). Similarly, incubation of human prostate tissue slides the *fimH* deletion mutant or WT in the presence of D-mannnose showed a striking bacterial adhesion impartment (**Figure 3c**, comparison to the WT UPEC in **Figure 2d**). This suggests that FimH is also the main adhesin utilized by UPEC to interact with prostate cells.

### FimH binds to the prostate-specific membrane receptor Ppap

Since FimH was proven to be vital for the invasion of prostate cells by UPEC, we next wondered which host receptors are involved in this interaction. Since the 2D model grown in the differentiation medium exhibited increased bacterial invasion, we opted to use this model moving forward. To date, four receptors have been described to directly interact with FimH in the bladder: UPK1A^31^, uromodulin (UMOD)^32^, integrin beta 1 (ITGB1) and integrin alpha 3 (ITGA3)^11^. In addition, desmoglein 2 (DSG2) has recently been described as a FimH receptor in kidney cells^33^. Given that UPK1A and UMOD are not expressed in the human prostate tissue, ITGB1 is very lowly expressed in prostate cells, and ITGA3 is only expressed by basal prostate cells (**Extended Data** Figure 7a)^34^, we hypothesised that FimH might not bind to any of these known receptors but rather to a prostate-specific receptor/s yet to be identified. To test this hypothesis, we used a previously described protocol to block FimH binding to Itgb1, and Itga3 using antibodies against those proteins^11^. Because we did not find any antibody available against the extracellular domain of Dsg2, we could not test whether Dsg2 blocking could affect bacterial invasion. CFU results showed no difference in UPEC invasion between cells blocked for Itgb1 or Itga3 compared to a non-blocked control in the prostate model (**Extended Data** Figure 7b). Of note, using the bladder cell line 5637 as control showed a strong reduction in bacterial invasion (**Extended Data** Figure 7b), as previously published^11^. This suggests that these two integrins are not necessary for FimH attachment to the luminal prostate cells.

To identify the potential FimH receptor in prostate cells, we first produced a recombinant version of the lectin domain (LD) of FimH (rFimH_LD_) from the same UTI89 strain used in our studies with a 3xFLAG-tag fused to the C-terminus^35^. First, to test whether the rFimH_LD_ could bind to the host cells, we incubated both prostate cells as well as the bladder epithelial cell line 5637 as a control with the recombinant protein for 1 hour, and visualised rFimH_LD_ by immunofluorescence staining with an anti-FLAG antibody. In 5637 bladder cells, rFimH_LD_ was able to bind and showed the expected pattern with a high signal in the intercellular junctions, where most ITGB1 and ITGA3 are located (**Figure 4a**). Of note, adding D-mannose to the incubation medium abrogated the binding of rFimH_LD_ to the cells (**Figure 4a**). Compared to bladder cells, rFimH_LD_ bound to the prostate cells with a specific signal not only in the cell-cell junctions but also on the apical side of the cells (**Figure 4b**). In addition, we observed that rFimH_LD_ bound preferentially to some cells while others would remain negative for the staining. This suggests that FimH receptor/s might be confined to a specific cell type. Moreover, incubation of rFimH_LD_ with human prostate tissue slides showed a clear colocalisation with the apical side of the prostate luminal cells (**Figure 4c**).

**Figure 4.**
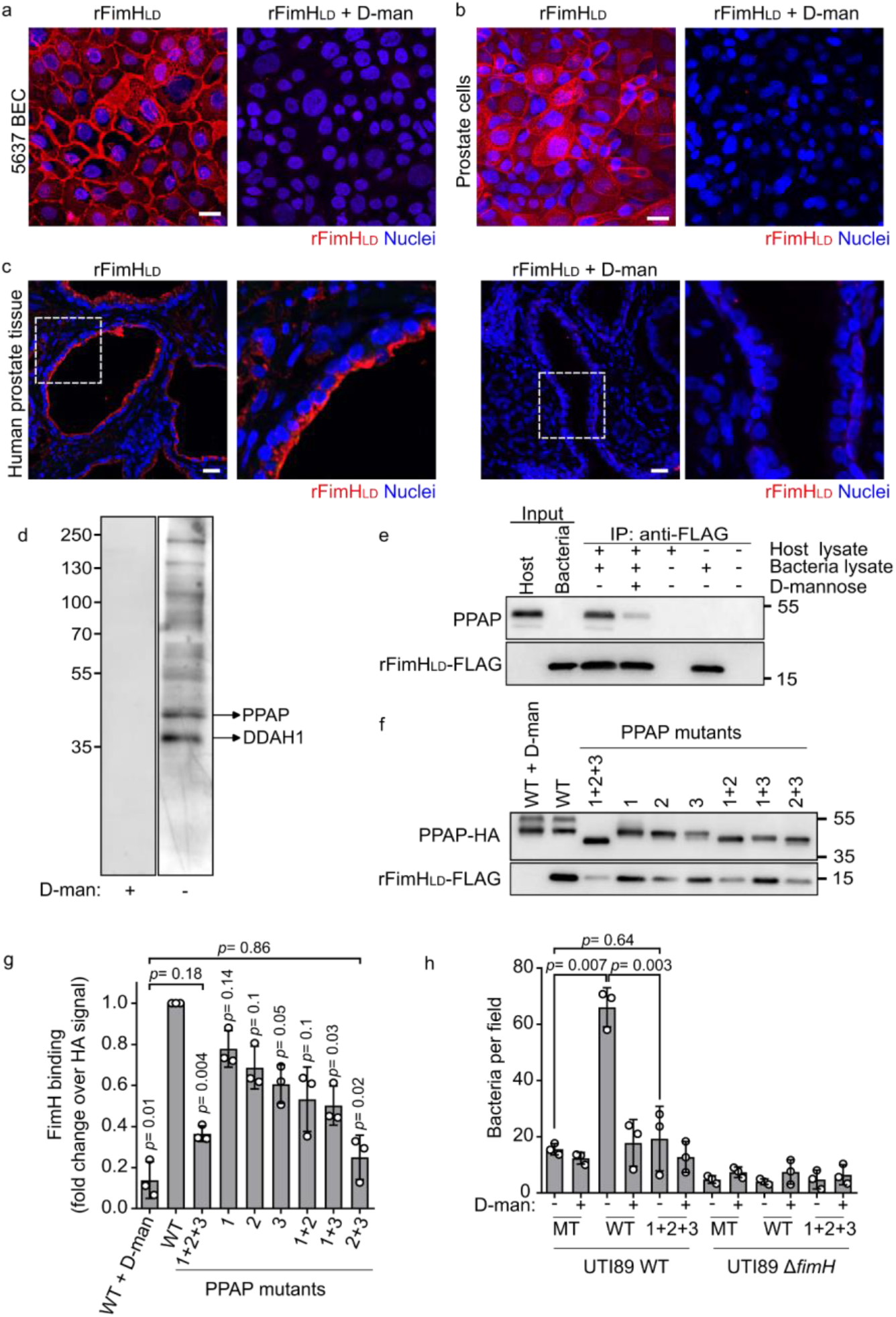
FimH binds to the prostate-specific membrane protein PPAP. a-c. Representative confocal microscopy images of recombinant FimHLD (rFimH_LD_) binding to the 5637 bladder cell line (**a**), organoid-based model grown with 10 nM DHT (**b**), and human prostate tissue (**c**). rFimH_LD_ signal (anti-FLAG antibody) is shown in red. Nuclei were counterstained using Hoechst 33342 (blue). Scale bar: 25 µm. (*n* = 3) **d.** Overlay assay: human prostate lysates enriched for membrane proteins were resolved in a SDS-PAGE, transferred to PVDF membrane and incubated with the rFimH_LD_- FLAG in the presence or absence of 2.5% of D-mannose. Prostate protein candidates were detected using an anti-FLAG antibody. The two most prominent bands were excised and identified as PPAP and DDAH1. **e.** Co-immunoprecipitation of PPAP using rFimH_LD_-FLAG as bait (*n* = 3). **f-g.** Representative blot (**f**) and quantification (**g**) of rFimH_LD_-FLAG binding to rPPAP-HA WT or mutants (mutant 1-position 94, mutant 2- position 220, and mutant 3- position 333; Fold change over HA signal; *n* = 3). **h.** Binding array showing UTI89 WT or UTI89 Δ*fimH* mutant binding to rPPAP-HA (WT and the triple mutant). Mock-transfected cell lysate was used as control (*n* = 3).

To identify potential FimH receptor/s in prostate cells, we used a similar approach previously applied to ITGA3 and ITGB1 as FimH receptors (far Western blot overlay assay)^11^. For that, host membrane proteins isolated from human prostate tissue were resolved by SDS-PAGE, transferred to a polyvinylidene fluoride (PVDF) membrane, blocked, and incubated with the rFimH_LD_. Incubation with an anti-FLAG antibody revealed several bands indicating rFimH_LD_ bound to several different host proteins (**Figure 4d**). Addition of D-mannose to the incubation abrogated any binding of rFimH_LD_ to the membrane as expected. The two most prominent bands bound to rFimH_LD_ had molecular weights of approximately 35 kDa and 45 kDa. Mass spectrometry analysis of the bands identified them as N(G), N(G)-dimethylarginine dimethylaminohydrolase 1 (DDAH1) and Prostatic acid phosphatase (PPAP), respectively. Because DDAH1 is a cytoplasmic protein, and PPAP is either secreted or a plasma membrane protein, we focused on PPAP as a potential FimH receptor.

The binding between rFimH_LD_ and PPAP was validated by immunoprecipitation using the rFimH_LD_ as bait (**Figure 4e**). Given that the binding of FimH adhesin to host proteins is mediated by the adhesion of the FimH lectin domain to high-mannose type N-glycans of glycoproteins, we tested whether mannosylated residues in PPAP were necessary for the binding of the adhesin to the host receptor. First, we generated a PPAP recombinant protein fused to an HA tag, which also served to validate PPAP binding to rFimH_LD_ (**Extended Data** Figure 8). PPAP has three N-glycosylated asparagines at positions 94, 220, and 333 that contain different numbers and forms of glycosyl units^36^. Hence, we mutated all three asparagines to alanines, either separately or in different combinations, and tested their respective contribution to the FimH binding. Single mutations showed a very small reduction in rFimH_LD_ binding (**Figure 4f-g**). However, a strong reduction was observed in the double mutant for positions 220 (position 2) and 333 (position 3; 0.25 ± 0.11 FC, *p* = 0.02). A similar effect was observed in the triple mutant (0.36 ± 0.04 FC, *p* = 0.004, **Figure 4f-g**), suggesting that glycosylation in position 93 (position 1) is not as crucial as glycosylation of positions 2 and 3. The addition of D-mannose to the reaction drastically reduced FimH capacity to bind to PPAP (**Figure 4f-g**) to a greater extent than the 2+3 or triple mutant, though the difference was not statistically significant (p = 0.86, *p*= 0,18, respectively). Moreover, to confirm PPAP binding to the full-length FimH adhesin expressed by UTI89 during infection, we performed a binding array assay with WT UTI89 bacteria. In this assay, rPPAP was bound to a glass microscopy slide and incubated with live bacteria, washed, and subsequently fixed. WT UTI89 bacteria were found to bind to the WT PPAP in higher numbers than to the triple mutant PPAP (**Figure 4h**). As expected, WT UTI89 bacteria in the presence of D-mannose or the FimH deletion mutant were not able to bind to either WT PPAP or the mutated PPAP (**Figure 4h**). Together, this data shows that UPEC FimH binds to the prostate receptor PPAP and that PPAP glycosylation is important for this interaction.

### Ppap is required for maximal UPEC invasion into prostate cells

Given that FimH assists bacterial invasion into the bladder cells^37^ and that FimH is also necessary for invasion into mouse prostate cells (**Figure 3a, b**), we hypothesised that the presence of Ppap on the cells would participate in the invasion process. Thus, to test if Ppap affects bacterial invasion, we generated two murine *Ppap* knock-out (KO) organoid lines using CRISPR/Cas9 technology as previously described^38^.

Sanger sequencing and FISH microscopy confirmed the successful knockout of the Ppap receptor in the murine organoid cells (**Extended Data** Figure 9a-c). Infection of the *Ppap* KO cells with UTI89 resulted in a significant decrease in bacterial invasion at 1hpi (9.6x for mouse #1 and 4.7x for mouse #2, *p*=0.005 and *p*= 0.02, respectively; **Figure 5a-b**), confirming the essential role of Ppap in achieving maximal UPEC invasion.

**Figure 5.**
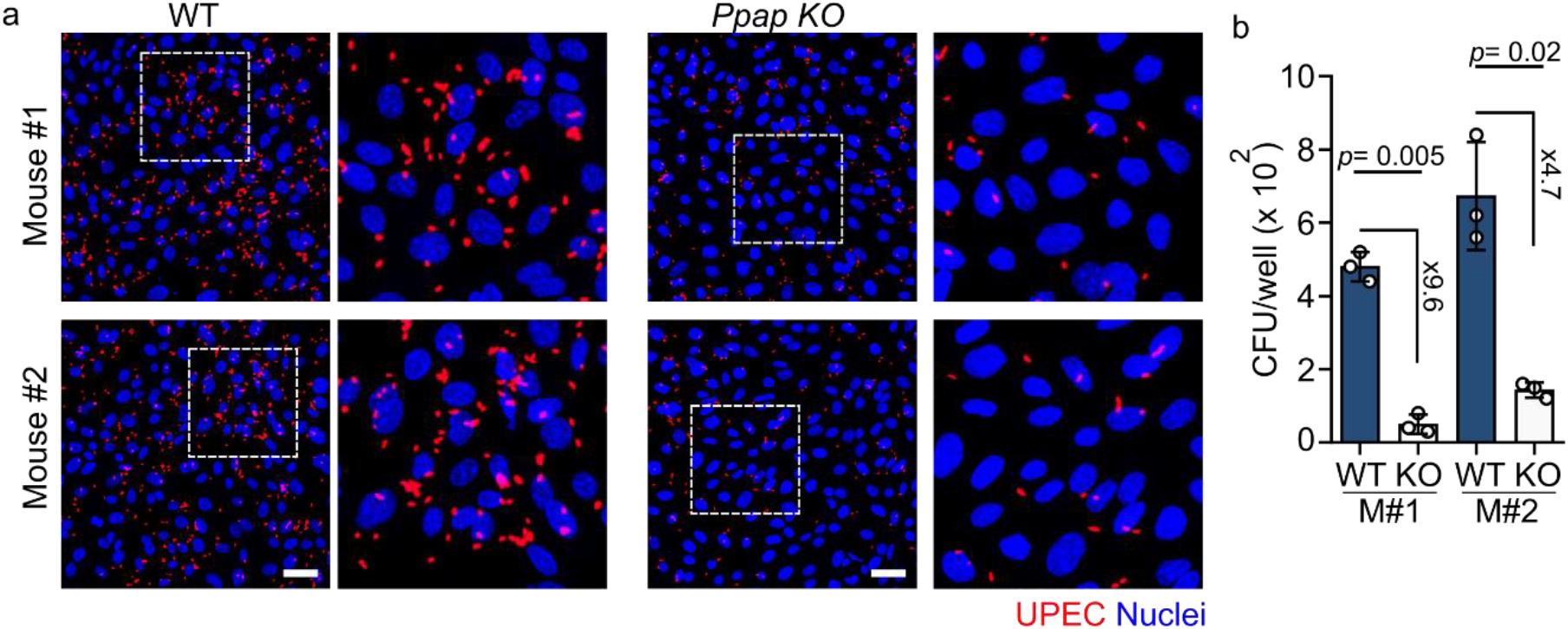
Ppap is necessary for maximal UPEC invasion into prostate cells. a-b. Representative confocal microscopy images (**a**) and CFU quantification (**b**) of two *Ppap* KO mouse organoid lines (grown in 10 nM DHT) infected with UTI89 WT (red) for 1h. Nuclei were counterstained using Hoechst 33342 (blue). Scale bar: 25 µm. *n* = 3.

### FimH colocalizes with PPAP in human prostate tissue

Having observed that FimH binds to PPAP *in vitro*, we next examined whether this binding could be also observed *ex vivo*. First, we tested whether live WT bacteria showed a preference for PPAP-positive cells in the tissue. To assess this, prostate tissue slides were incubated with live WT UTI89 bacteria (with or without D-mannose) and subsequently stained for PPAP. Confocal microscopy analysis showed that UTI89 was predominantly found close to PPAP-positive cells (**Figure 6a**). Validating our *in-vitro* data, the Δ*fimH* mutant strain or the WT supplemented with D-mannose was not able to bind to the prostate tissue (**Figure 6a**). In addition, following the same experimental approach as in **Figure 4a**, incubation of the tissue slide with the rFimH_LD_ protein showed a clear colocalisation with the PPAP signal in the luminal prostate cells. Overall, these results demonstrate that UPEC binds to prostate luminal cells expressing PPAP on their surface via FimH.

**Figure 6.**
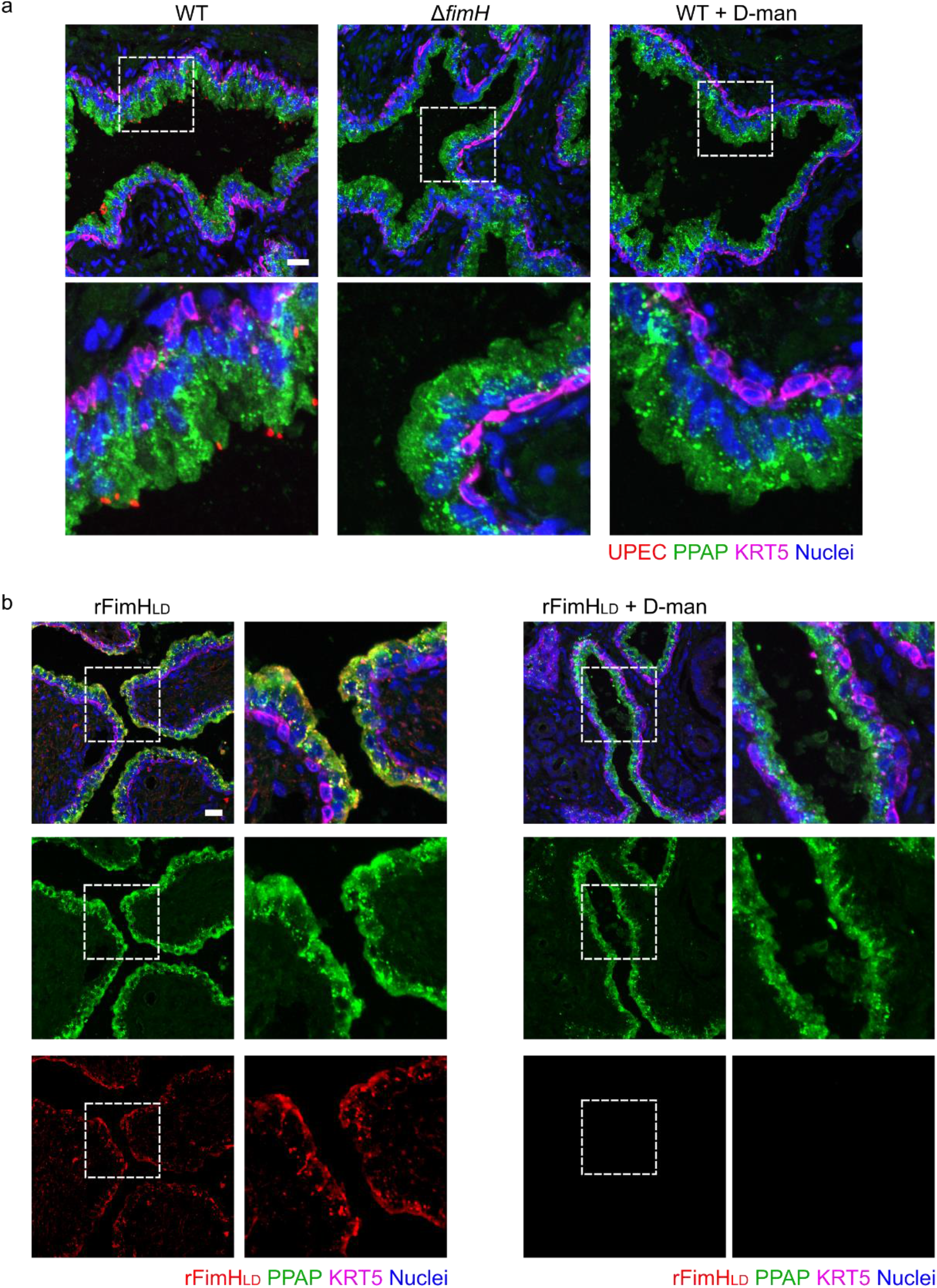
rFimH_LD_ and UPEC colocalizes with PPAP positive cells in human prostate tissue. **a.** Representative confocal microscopy images showing the binding of UTI89 WT, Δ*fimH* mutant, and WT in the presence of 2.5% D-mannose to human prostate tissue. Bacteria are shown in red, PPAP in green and KRT5 in magenta. **b.** Representative confocal microscopy images of rFimH_LD_ (red) binding to human prostate tissue and stained for PPAP (green) and KRT5 (magenta). Nuclei were counterstained using Hoechst 33342 (blue). Scale bar: 25 µm. Data are representative results from 2 donors and 2 independent experiments.

## DISCUSSION

UPEC is a versatile bacterial pathogen that can infect various organs in the genitourinary tract, using distinct strategies for each tissue. In this work, we established an adult stem cell organoid-based model of the mouse prostate to study the main hallmarks of UPEC infection in the prostate epithelium, an understudied tissue susceptible to UPEC infection. Using this model, we show that UPEC is able to adhere to, invade, and replicate within prostate luminal cells and that this is mediated by the binding of FimH to the glycosylated prostate-specific membrane receptor PPAP (**Figure 7**).

**Figure 7.**
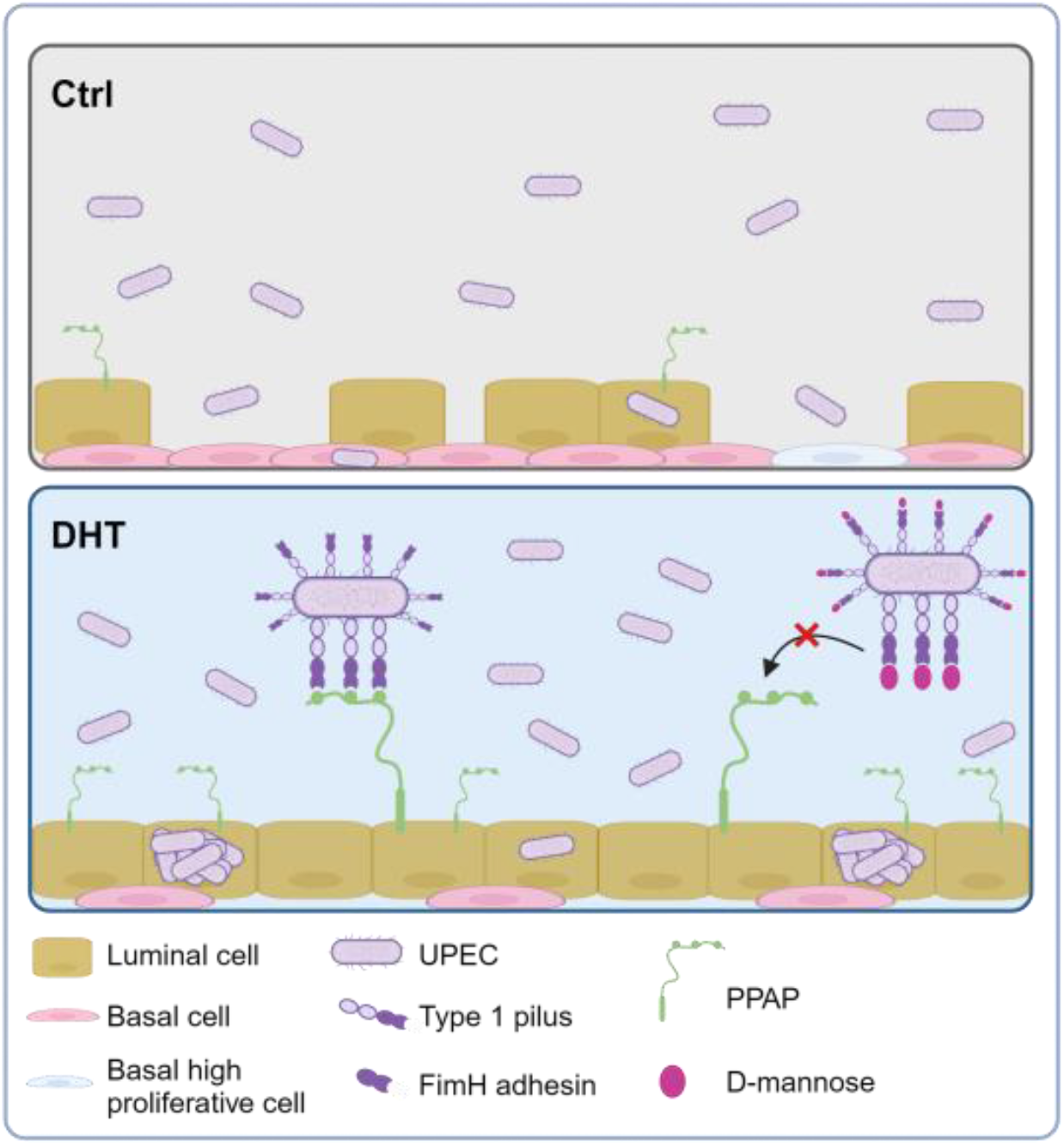
Model depicting the role of the PPAP protein in UPEC infection of prostate cells. In the organoid model exposed to 10 nM DHT, cells differentiate into luminal cells, while in the absence of DHT (control), they remain as basal cells. UPEC targets prostate luminal cells by using FimH to bind to the PPAP receptor on their surface, facilitating bacterial invasion. D-mannose can inhibit this process by blocking FimH binding to PPAP, thereby preventing invasion.

Recent advances in organoid technology have significantly improved infection biology by enabling more accurate *in-vitro* studies of pathogens in primary cells^39–42^. These primary cells can differentiate into the various epithelial cell types found in tissues, given the appropriate culture conditions. This is crucial for studying infectious diseases, as pathogens interact differently with distinct host cell sub-types. Therefore, ensuring that the *in-vitro* model accurately mimics epithelial heterogeneity is essential for effective research. Here, using mouse prostate adult stem cell organoids, we have developed a differentiated model that mimics the cell composition of the prostate epithelium, with the exception of the rare neuroendocrine cells. Our data shows that in 3D, organoids have a higher proportion of undifferentiated basal stem cell-like cells, while cells seeded on a 2D surface tend to differentiate more. This is also consistent with our previous study in gastric organoids, where 2D organoid-based model exhibited higher cell differentiation towards pit cells than 3D organoids under the same culture conditions^38^. This difference is likely due to the distinct physical cues that cells experience in 2D versus 3D environments. Furthermore, in our model, supplementing our culture medium with a ten-fold higher concentration of DHT induced higher cell differentiation of the basal stem-cell-like cells into luminal prostate cells, similar to the cell proportions in real tissue^23,25^. In this context, previous work on mouse organoids has shown that androgen deprivation decreases the number of luminal markers and that upon androgen signal restoration, luminal cells can restore the prostate epithelium^23,26^. Conversely, removing DHT directed cell differentiation towards a basal-like phenotype. Such a model, mainly consistent of basal stem cell-like cells, could hold additional benefits in studying, e.g. prostate basal cell carcinomas or the response of basal cells to specific stimuli (e.g. hormones).

In our study, UPEC exhibits a clear preference for luminal cells over basal stem cell-like cells. Previous studies have already described bacterial invasion and discrete replication in human prostate-transformed cell lines^14,43–45^. However, cell differentiation was not addressed in those studies. Despite the lack of *in vivo* studies exploring bacterial invasion in the prostate, our observation is consistent with findings in bladder infections, where bacteria replicate and form intracellular bacterial communities more efficiently in superficial umbrella cells than in intermediate cells. While the adhesion to differentiated umbrella cells could be explained by the presence of specific host receptors (e.g., uroplakin 1a)^31^, the higher intracellular replication rate in those cells is not well understood as of yet. Different hypotheses, such as the physical space in the host cell cytoplasm or the distribution of actin filaments, have been proposed^46,47^, but so far, neither has been proven. Nonetheless, bacterial invasion into bladder cells allows the pathogens to escape the immune response, antibiotic treatment, and elimination via micturition. Although the prostate gland is not void as frequently as the bladder, it seems plausible that invading the prostate luminal cells would benefit the bacteria in a similar manner.

Several studies have shown that testosterone increases susceptibility to and worsens the outcome of bladder and kidney infections, majorly due to its immunosuppressive effect in immune cells^27,48–51^. However, none of those studies looked at the effect of testosterone on the epithelial cells. In our model, adding DHT increased cell differentiation towards UPEC target cells, thus facilitating infection. Notably, DHT had no effect on UPEC growth, and it was removed from the medium during the short incubation of bacteria with cells to eliminate any potential influence of the hormone on UPEC. Other studies have reported reduced UPEC invasion and replication in prostate cell lines following treatment with DHT. However, the concentrations used in those studies were significantly higher—20 and 40 μg/ml (equivalent to 68.8 μM and 137.6 μM, respectively)^50^ or 5 and 20 μg/ml (equivalent to 17.2 μM and 68.8 μM, respectively)^45^—compared to our more physiological level of 10 nM^52^.

Bacterial adhesion to the host epithelial cell is a critical step in infection and a prerequisite to invasion. UPEC possesses a versatile range of adhesins or pili, including the type 1 pili, the pyelonephritis-associated pilus (Pap) and S pili, the Dr adhesins and others^52^, allowing the bacteria to successfully colonize the different niches. This is not trivial for the bacteria: while UPEC uses the PapG adhesins on P pili to bind to galactose-rich membrane glycolipids (e.g., globotetraosylceramides or globotriaosylceramides) on kidney cells^53^, it uses the type 1 pili to bind to mannosylated residues on the surface of bladder urothelial cells^37^ and vaginal epithelial cells (although other adhesins also contribute to colonization of female genital tract)^54,55^. Our data show that the FimH adhesin at the tip of the type 1 pili is the main adhesin responsible for UPEC binding to prostate cells. However, a small proportion of FimH mutant bacteria was still able to adhere to prostate cells, suggesting the potential presence of other adhesin/s or other surface structures contributing to it. FimH binds to different host receptors in the bladder, including UPK1a (present in umbrella cells)^31^, integrins ITGA3 and ITGB1 (present in umbrella and intermediate cells)^11^, and UMOD (secreted by kidney cells to the urine)^32^. Since these receptors are either absent from the prostate or present to a lower extent, it raised the question of whether there are other prostate-specific receptors for FimH. The binding of the recombinant FimH to the differentiated model showed a very distinct binding pattern with a spread signal at the apical side of the host cell, in contrast to the basolateral signal observed in bladder cells. Moreover, small, punctate, round structures were observed on the central apical side, which could suggest binding to prostatic extracellular vesicles or prostasomes. Our results indicate that the PPAP protein is a novel FimH receptor in the prostate epithelium. PPAP is a non-specific tyrosine phosphatase exclusively expressed by luminal prostate cells that dephosphorylates a diverse number of substrates under acidic conditions^56–59^. This, in turn, helps sperm motility, although the exact mechanism is not well understood^60^. It also inhibits prostate cancer through dephosphorylation of ERBB2 and deactivation of MAPK-mediated signalling^58^. Both human and murine PPAP are present in two isoforms that are produced through alternative splicing: isoform 1 is secreted, and isoform 2 is a type-I transmembrane protein^61^. Because only isoform 2 could function as a receptor in the cells, we focused on this isoform. Nevertheless, the secreted isoform might also act as a soluble FimH receptor, similar to uromodulin. Our knockout organoid lines showed that Ppap is necessary to achieve the maximum UPEC invasion in differentiated cells. However, a small proportion of intracellular UPEC was found in the KO cells, suggesting the potential presence of other host receptor/s. This aligns with FimH binding observed in other tissues, where FimH can engage with different receptors within the same tissue.

All the FimH receptors described to date are heavily mannosylated glycoproteins. Similarly, human PPAP contains three N-glycosylation motifs (Asn94, Asn220, and Asn333)^36,62^ that could serve as anchors for the FimH lectin binding pocket. Of note, since both isoforms (secreted and membranal) exhibit the same glycosylation pattern, this enforces the hypothesis that soluble PPAP could serve as a soluble factor for FimH. The three positions have different numbers of glycosyl units, with the Asn333 being the most glycosylated or heavy chained^36^. We hypothesised that removing PPAP glycosylation by substituting asparagine with alanine at each position would affect the ability of FimH to bind to PPAP. Our results showed that while two glycosylated positions (Asn220 and Asn333) were essential for FimH binding, one of them (Asn94) appeared less important.

In our study, both the recombinant FimH protein and live UPEC bacteria demonstrated a clear preference for human luminal cells positive for PPAP protein in human prostate tissue. To our knowledge, this is the first time that UPEC interaction has been observed in *ex vivo* human prostate tissue. Although the colocalisation of the PPAP signal with UPEC or rFimH_LD_ in the tissue does not directly confirm binding, it strongly suggests that our *in-vitro* data reflect a more physiologically relevant context, thereby reinforcing the significance of our findings.

The present work underscores the significance of type 1 pili and the mannosylated epithelial receptor PPAP in UPEC infection of prostate cells. Since FimH is already targeted by small-molecule inhibitors and vaccines in bladder infection patients^63,64^, these therapeutics could also be effective in preventing or treating bacterial prostatitis.

## METHODS

### Reagent and resource sharing

Further information and requests for resources should be directed to C.A.

### Organoids generation

Mouse prostate organoids were generated from WT male C57BL/6 mice as described before^24,26^ with minor modifications. Briefly, prostate tissue was minced into small pieces and digested in 5 mg mL-1 collagenase (C9891-100MG, Sigma Aldrich) with 10 µM RHOKi (Y-27632; M1817, AbMole) for 1 h at 37°C on a shaking platform. Tissue fragments were then washed once with Advance Dulbecco’s modified Eagle medium/F12 (12634028, Thermo Fisher Scientific) supplemented with 10 mmol L−1 HEPES (15630056, Thermo Fisher Scientific), 1x GlutaMAX (35050-038, Thermo Fisher Scientific; adDMEM/F12+/+) and centrifuged at 150 × *g* for 5 min at 4°C. The cell pellet was resuspended in TrypLE Express Enzyme (12605028, Gibco) supplemented with 10 µM RHOKi (Y-27632, Sigma-Aldrich) for 15 min at 37°C. The cell suspension was then washed, filtered with a 100 µm strainer (43-10100, pluriSelect), and centrifuged at 350 × *g* for 5 min at 4°C. Cells were seeded in 50 µL drops of Matrigel (356231, Corning). Culture medium contained: Advance Dulbecco’s modified Eagle medium/F12 (12634028, Thermo Fisher Scientific), 10 mmol L−1 HEPES (15630056, Thermo Fisher Scientific), 1x GlutaMAX (35050-038, Thermo Fisher Scientific), 1xB27 (12587010, Thermo Fisher Scientific), 1 mM N-Acetylcysteine (A9165-25G, Sigma-Aldrich), 10% Noggin conditioned medium, 10% R-spondin 1 conditioned medium, 50 ng mL−1 EGF (AF-100-15, Peprotech), 1 nM dihydrotestosterone (5α-Androstan-17β-ol-3-one, DHT; A8380, Sigma-Aldrich), 0.2 µM TGFβi (A-83-01, Tocris), and 100 ng mL−1 Primocin (Ant-pm-1, Invivogen). For propagation, organoids were split weekly in a ratio 1:4. To adhere to the 3R principles (Replacement, Reduction, Refinement), we utilised only leftover material from mice previously used for other experiments. No additional animals were sacrificed for this study.

### Organoid-based model seeding

One week old mouse prostate organoids were used to seed the organoid-based model. Organoids were collected and mechanically disrupted. Organoid fragments were washed with adDMEM/F12+/+ and centrifuged at 450×g for 5 min. Organoid fragments were then resuspended in TrypLE Express (12605028, Gibco) and incubated at 37 °C for 15 min. Single-cell suspension was washed and centrifuged at 450 × *g* for 5 min. Single cells were counted and seeded at 12,500 cells per well cells/well in 48-well plate (833923, Sarstedt) using the organoid culture medium without Primocin and let to grow for a week. RHOKi (Y-27632; 10 μM, M1817, AbMole) was added at the seeding time. To induce cell differentiation, 10nM DHT was added to the medium on the day of splitting or seeding and kept throughout the experiment. Ethanol was used as a vehicle control in the control condition.

### Bacterial culture and infection

UPEC strain UTI89 WT expressing mCherry or GFP were obtained by transforming the bacteria with the plasmid pFPV or pXG10, respectively. Chromosomal deletion of *fimH* in UPEC strain UTI89 was achieved by λ Red recombineering^65^. Briefly, a DNA fragment including the cat cassette flanked by FRT sites and 60 bp homologous regions to the 5′ and 3′end of fimH was PCR-amplified from pKD3 using the primers fimH::cat_npt_ff(ATGAAACGAGTTATTACCCTGTTTGCTGTACTGCTGATGGGCTGGGTGTAGGCTGGAGCTGCTTC) and fimH::cat_npt_rev (TTGATAAACAAAAGTCACGCCAATAATCGATTGCA CATTCCCTGCCATATGAATATCCTCCTTAGTTCC). UPEC was then transformed with 250 ng of the obtained PCR product, and mutants were selected on LB agar plates supplemented with 12.5 µg/ml chloramphenicol. The deletion of *fimH* was verified by PCR and the mutant was cured from pKD46 by incubation at 42 °C.

For in vitro infection, bacteria were grown in LB broth overnight at 37°C under static conditions. A 24-hour culture was inoculated with OD600 0.05 from the overnight culture and grown under the same conditions. Bacteria were harvested by centrifugation and resuspended in organoid culture medium. Bacteria were then diluted to the desired MOI in the organoid culture medium and added to the wells. D-mannose (M6020, Sigma Aldrich) at a final concentration of 2.5% was added to the medium simultaneously with the bacteria when indicated. After 1 h incubation, the unbound bacteria were washed, and the culture medium was exchanged with fresh medium supplemented with 50 µg mL-1 gentamicin (G1272, Sigma Aldrich). After 30 min, the medium was changed to a lower concentration of gentamicin (10 µg mL-1; G1272, Sigma Aldrich) and left until analysis. At the indicated time, the cells were washed with PBS and collected for further analysis. For OD600 measurements over time (growth curves), 24-hour cultures were inoculated in a 96-well plate to an initial OD600 of 0.01 in the presence of 10nM DHT or ethanol (vehicle).

### CFUs

A CFU (Colony Forming Units) assay was used to quantify intracellular bacteria. At the indicated time, organoid-infected cells were washed with PBS and lysed with 0.1% Triton X-100 (T9284, Sigma-Aldrich) in PBS. Cell lysates were then serially diluted and plated on LB agar plates and grown overnight.

For the antibody-blocking, experiments were done as previously published^11^. Briefly, cells were pre-incubated with purified monoclonal antibodies for 30 min before adding bacteria. The final concentration of each antibody was 1.5 μg/well. The following antibodies were used: β1 integrin (6S6; sc-53711, Santa Cruz Biothecnology) and α3 integrin (P1B5; sc-13545, Santa Cruz Biothecnology). PBS was used as control in the “No Ab” condition.

### Staining and immunofluorescence

Cells seeded on a µ-Slide 18 Wells (81816, Ibidi) were washed with PBS and fixed with 4% paraformaldehyde (PFA) for 15 min at room temperature, permeabilised with 0.5% Triton X-100 (T9284, Sigma-Aldrich) in PBS for 15 min and then blocked with blocking buffer (1% BSA in PBS) for 30 min at room temperature. Primary antibodies were diluted in blocking buffer and incubated overnight at 4°C followed by 1 h at room temperature. Anti-Cytokeratin 5 (ab52635, Abcam), anti-Cytokeratin 5 Alexa Fluor® 647 antibody (ab193895, Abcam), anti-CD24a (10600-1-AP, Proteintech), and anti-ACPP (PPAP, LS-C292593, LSBio), anti-KRT18 (ab133263, Abcam) antibodies were used at 1:100. Anti-ZO-1 Polyclonal antibody (21773-1-AP, Proteintech) was used at 1:750. Cells were washed with PBS and incubated with the secondary antibodies anti-Rabbit IgG Alexa Fluor™ 488 (A21441, Invitrogen) or anti-Mouse IgG Alexa Fluor™ 594 (A21201, Invitrogen) at a dilution of 1:500. To stain F-actin, cells were stained with Flash Phalloidin™ Green 488 (424201, Biolegend) or Red 594 (424203, Biolegend) at 1:250 dilution for 1 h. For differential staining of intracellular versus extracellular bacteria, after fixation with PFA 4%, the permeabilisation step was omitted. After blocking with 1% BSA, extracellular bacteria were stained with anti-*E. coli* LPS (ab35654, Abcam), and the anti-Mouse IgG Alexa Fluor™ 594 (A21201, Invitrogen) at a dilution of 1:500. Nuclei were counterstained with Hoechst 33342 (1:5,000; H3570, Life Technologies).

To stain and image the human prostate tissue, tissue explants were obtained from patients undergoing surgery at the University Hospital of Würzburg. This study was approved by the ethical committee of the University of Würzburg (Approval 168/22), and informed consent was obtained from all the donors. Tissue pieces were fixed with 4% PFA, dehydrated overnight with 30% sucrose and then embedded in O.C.T. Tissue slides were then re-hydrated with PBS for 10 min, and immunostaining was performed as described for the Ibidi slides.

Confocal microscopy images, shown as maximum projected Z-stack images, were acquired with a Leica SP5 laser scanning confocal microscope or with Leica Stellaris 5 confocal microscope and LAS AF Lite software (Leica Microsystems). The images were analysed with ImageJ.

### Bacterial incubation of human prostate tissue slides

Prostate tissue slides were obtained as described above. After re-hydration, slides were incubated with UPEC UTI89 WT or Δ*fimH* (OD600 0.02) for 30 min on a shaking platform in the presence or absence of D-mannose (2.5%, M6020, Sigma Aldrich). The slides were then washed with PBS, fixed with PFA 4%, and further processed for immunofluorescence as described above.

### Data mining from “The Human Protein Atlas”

IHC images of human prostate tissue samples stained for UPK1A, UMOD, ITGB1, ITGA3, and DSG2 were obtained by datamining the human protein atlas database (www.proteinatlas.org)^34^.

### TEER measurement

ECIS 8-well transfilter Array connected to ECIS® Z-Theta (Applied Biophysics) was used to measure the TEER of the organoid-based model. Cells were seeded as described above on a 8-well PET slide with 40 electrodes per well (8W10E+, Applied Biophysics). TEER values were measured at 400 Hz for 48 h (5-7 days after seeding).

The TEER of a blank well (only culture medium) was used as control and subtracted from all TEER values.

### rFimH_LD_ production and far western blot overlay assays

The lectin domain of *fimH* (1-156 amino acids) of UTI89 was amplified from genomic DNA using the forward primer 5’- CCCCGAATTCAGGAGGAGATTGTAATGAAACGTGTTATTACCCTG-3’ and the reverse primer 5’- GCGCGGATCCTTACTTATCGTCGTCATCCTTGTAGTCGCCGCCAGTAGGCACCAC-3’, cloned into the EcoRI-BamHI sites of plasmid pQE-30 and transformed into *E. coli* T4 Express competent cells (C3037I, New England BioLabs). Expression of the fimH-FLAG was induced with 1 mM IPTG (2316.3, Carl Roth) for 5 h. Bacterial cells were harvested and resuspended in cold 20 mM Tris/20% sucrose solution along with the protease inhibitor cocktail (05892791001, Sigma Aldrich) and 200 µM PMSF. The cells were then sonicated and incubated with 75 µg mL-1 of lysozyme for 30 min. Cell debris was removed by centrifugation at 15,000 × *g* at 4°C for 20 minutes. Supernatant was filtered twice through a 0.45 µm filter to remove smaller debris. Proteins were then concentrated and washed with 20 mM Tris using a 10 kDa molecular weight cutoff centrifugal filter (88527, Thermo Scientific). The presence of rFimH_LD_ protein was confirmed by western blot analysis using the antibodies anti-FimH (CSB- PA362349ZA01ENV, Bio Trend, 1:1,000) and anti-FLAG (1:1,000; 66008-4-Ig, Proteintech).

Far western blot overlay assays were done as described before^11^ with minor modifications. Human prostate tissue from four patients was lysed using the Proteoextract Native Membrane Extraction kit (Sigma Aldrich, 444810-1KIT) to enrich for membrane proteins. Pooled protein lysates were then separated in a 10 % SDS PAGE Gel. The proteins were then transferred on a PVDF transfer membrane and blocked with 1 % BSA/1 % milk in TBS-T (TBS 0.1% Tween-20, pH-7.4) for 20 min. The membrane was incubated with the bacterial lysate containing the rFimH_LD_ at a final concentration of 100 µg mL-1, alone or in the presence of 2.5 % D-Mannose (M6020, Sigma Aldrich) diluted in the blocking buffer for 90 min. After TBS-T washes, the membrane was incubated with an anti-FLAG antibody (66008-4-Ig, Proteintech, 1:1,000) for 1 h, washed, and incubated with the secondary anti-mouse IgG-HRP conjugate (GENA931, Sigma Aldrich, 1:10,000). Signals were detected using SuperSignal West Dura Extended Duration Substrate (Pierce, 34075) using an ImageQuant LAS 4000 CCD camera (GE Healthcare). Host protein bands that bound to rFimH_LD_, were excised from the duplicate SDS page gel that was stained with Coomassie Blue and identified by mass spectrometry at the Mass Spectrometry Facility of the Rudolf Virchow Center for Integrative and Translational B (University of Würzburg).

### rPPAP protein expression

The PPAP isoform 2 (membrane protein) protein tagged with HA in the C terminus was expressed using a commercially available plasmid from Sino Biologicals (HG10959-CY, Sino Biologicals). The plasmid was transfected into HeLa 229 cells using X-tremeGENE™ HP DNA Transfection Reagent (6366236001, Merck) in a 1:2 ratio. Cells were collected 48 h post-transfection in lysis buffer containing 1% NP-40, 5% glycerol, 25 mM Tris, 150 mM NaCl, 1 mM EDTA and protease inhibitor cocktail (05892791001, Sigma Aldrich). Cell lysis was completed by incubation on ice for 1 h, followed by centrifugation at 15,000 × *g* for 1 h. Pellets were discarded, and the PPAP expression was examined by Western blot using the anti-HA Polyclonal Antibody (51064-2-AP, Proteintech, 1:5,000) and the secondary anti-rabbit IgG-HRP conjugate (GENA934, Sigma Aldrich, 1:10,000).

### Co-immunoprecipitation

Co-Immunoprecipitations were performed using the Pierce™ Anti-DYKDDDDK Magnetic Agarose (A36797, Thermo Fisher Scientific) or the Pierce™ anti-HA magnetic beads (13464229, Thermo Scientific) to pull down rFimH-FLAG or PPAP- HA-binding partners, respectively, and following the manufacturer’s instructions. Briefly, for the co-immunoprecipitation with the Pierce™ Anti-DYKDDDDK Magnetic Agarose (A36797, Thermo Fisher Scientific), bacteria protein lysates were obtained as described before and incubated with the beads (100 µl lysate, 30 µl beads) for 30 min at 37C. Beads were then washed with wash buffer (20 mM Tris-HCl, 150 mM NaCl, pH-7.2) and incubated with 200 µl of the human prostate protein lysates for 30 min at 37C. Beads were then washed with wash buffer, eluted by boiling in 50 µl of Laemmli buffer and analysed by Western blot assay. An anti-PPAP antibody (HPA063916, Atlas Antibodies, 1:300) was used in a Western Blot assay to validate the co-immunoprecipitation of PPAP from the human prostate lysate with the rFimH_LD_- FLAG beads.

For co-immunoprecipitation using the Pierce™ anti-HA magnetic beads (13464229, Thermo Scientific), PPAP-expressing HeLa 229 cells (and mock controls) lysates were incubated with the beads (150 µl lysate, 20 µl beads) for 30 min at room temperature. Beads were then washed with wash buffer (TBS-T, 0.05% Tween 20) and incubated with 40 µl of bacterial lysate for 30 min at room temperature. Beads were then washed with wash buffer, eluted by boiling in 50 µl of Laemmli buffer and analysed by Western blot assay. For D-mannose blocking experiments, D-mannose (M6020, Sigma Aldrich) to a final concentration of 2.5 % was added to the bacterial lysate.

### rFimH_LD_ binding assays

To confirm that the rFimH_LD_ could bind to host cells, we first used the 5637 cell line (HTB-9, ATCC) as control. 5637 cells were cultured in RPMI 1640 GlutaMAX (Thermo Fischer, 72400047). A day before the experiment, 18,000 cells/well were seeded on a µ-Slide 18 Wells (81816, Ibidi). Next day, fresh medium containing 10% of the bacterial lysate (rFimH_LD_) was incubated with the cells for 1 h at 37C. Then, cells were washed, fixed with 4% PFA, blocked with 1% BSA and stained with an anti-FLAG antibody (1:1,000; 66008-4-Ig, Proteintech) for 1 h at room temperature. Then cells were incubated with an anti-Mouse IgG Alexa Fluor™ 594 (1:500; A21201, Invitrogen) for 1 h at room temperature. D-mannose (M6020, Sigma Aldrich) at a final concentration of 2.5% was added to the medium at the same time as the bacterial lysate when indicated. The same procedure was performed on the organoid-based model (DHT condition). In this case, organoid cells were seeded as described above. Nuclei were counterstained with Hoechst 33342 (1:5,000; H3570, Life Technologies).

For the incubation of the human prostate tissue slides, O.C.T-embedded tissue slides were re-hydrated for 10 min in PBS at room temperature, blocked with 1% BSA and incubated with blocking buffer containing a 10% protein lysate in the presence or absence of D-mannose (2.5%, M6020, Sigma Aldrich) for 1h at room temperature. Then, slides were washed and fixed in PFA 4%. Staining of rFimH_LD_-FLAG was done as described for the Ibidi slides above.

### Site-directed mutagenesis

Mutants of *ACP3* (*PPAP*) putative binding sites were generated by site-directed mutagenesis. A pair of complementary primers carrying the desired point mutation at positions 94 and pos. 220 (94:5′- AAGAGATATAGAAAATTCTTGGCTGAGTCCTATAAACATGAACAG-3′ and 5′- CTGTTCATGTTTATAGGACTCAGCCAAGAATTTTCTATATCTCTT-3′, 222: 5′-TATATTGTGAGAGTGTTCACGC-3′ and 5′-CAGGAGGGTAAAGTGAAAGC-3′) were used to introduce a point mutation that mutated the asparagine to alanine by PCR. This approach was unsuitable for generating the same point mutation for position 333 due to the high GC content of the sequence around the position. Hence, a DNA sequence containing the point mutation in position 333 was obtained from IDT (TGCGCATGACACTACTGTGAGTGGCCTACAGATGGCGC TAGATGTTTACAACGGACTCCTTCCTCCCTATGCTTCTTGCCACTTGACGGAATT GTACTTTGAGAAGGGGGAGTACTTTGTGGAGATGTACTATCGGGCTGAGACGC AGCACGAGCCGTATCCCCTCATGCTACCTGGCTGCAGCCCCAGCTGTCCTCTG GAGAGGTTTGCTGAGCTGGTTGGCCCTGTGATCCCTCAAGACTGGTCCACGGA GTGTATGACCACAAACAGCCATCAAGTTCTAAAGGTCATCTTTGCTGTTGCCTTT TGCCTGATATCTGCTGTCCTAATGGTACTACTGTTTATCCACATTCGCCGTGGA CTCTGCTGGCAGAGAGAATCCTATGGGAACATCGGGGGTGGAGGCTCTTATCC TTACGACGTGCCTGACTACGCCTAAACTCGAGTCTAGA; Integrated DNA

Technologies). The sequence was cloned into the ACP3 plasmids (WT or mutants 93, 222 or both) between the XbaI and FspI restriction sites. Correct plasmid DNA sequences were confirmed by Sanger sequencing.

### Bacterial binding to immobilized PPAP assays

For bacterial adhesion assays on immobilised PPAP-HA WT and mutants, cell lysates from PPAP-HA expressing HeLa cells were immobilized on PDL coated glass slides (J2800AMNZ, Epredia™) using a modified protocol already described^66^. Briefly, the Grace Bio-Labs ProPlate microarray system for 64 wells (GBL246865; Grace Bio- Labs) was used on PDL coated glass slides. Slides were incubated with 12.5 µg mL- 1 of anti-HA (51064-2-AP, Proteintech), or anti-LPS (ab35654, Abcam) antibody in array buffer (3% BSA in PBS) for 2 h at room temperature. Afterwards, wells were washed with array buffer and incubated with 50 µL of cell lysate overnight at 4°C. Wells were then washed with array buffer before the addition of 50 µL (OD600 0.05) of UPEC strain UTI89 WT or ΔfimH mutant, in the presence of absence of D-mannose (2.5%, M6020, Sigma Aldrich). After incubation with bacteria, slides were washed, fixed with 4% PFA and imaged using the ECHO Revolve microscope. Bacterial numbers attached to the slide were in four different images per experiment and condition using ImageJ software.

### Single-cell RNA sequencing and Data analysis

To generate single-cell suspensions, organoids and organoid-based model cells were washed with PBS, and dissociated with TrypLE Express Enzyme (12605028, Gibco) for 15 min at 37°C. Samples were washed with adDMEM/F12+/+, centrifuged at 400 × *g* for 5 min and resuspended in PBS with 0.1% BSA. Cell viavility was measured with tripan blue. Only samples with over 90% viability were used. Cells from two mice (grown in control and DHT medium) were multiplexed in two pools (2D and 3D organoids) using CellPlex technology (10x Genomics). Samples were resuspended in CellPlex Tag Buffer and incubated with individual CellPlex oligonucleotide tags for 15 min at room temperature. After labelling, cells were washed twice with PBS containing 0.04% BSA and pooled together in equal proportions. Cell concentration and viability were determined using an automated cell counter (Countess II, Thermo Fisher Scientific).

The pooled cells were subsequently loaded onto a Chromium Controller (10x Genomics) to generate Gel Bead-In-Emulsions (GEMs). Libraries were quantified using a Qubit dsDNA HS Assay Kit (Thermo Fisher Scientific) and their size distribution was assessed using an Agilent 2100 Bioanalyzer. Sequencing libraries were then loaded onto an Illumina NovaSeq 6000 platform using a S2 flow cell with 100 cycles. Raw sequencing data was demultiplexed, quality checked and subsequently aligned and quantified using the Cellranger software suite (10x Genomics, v7.0.1) against the mm10 (Ensembl98) mouse genome assembly as reference.

Obtained count matrices of 3D organoids and 2D models were loaded into R (v4.3.0) and analysed analogously using the Seurat R package (v4.3.0). Cells were quality filtered based on standard quality metrics, including the number of detected transcripts per cell (>10000 2D, >5000 3D organoids), the number of detected genes per cell (> 200 < 9000 2D, >200 <8000 organoids), and mitochondrial count fraction (<10%). Raw counts were normalised and log-transformed (NormalizeData, default settings). Highly variable genes (HVGs) were identified (FindVariableFeatures, using nfeatures = 3000) and scaled (ScaleData, default settings). Principal component analysis (PCA) was performed based on HVGs (RunPCA, default settings). The uniform manifold approximation and projection (UMAP) algorithm was used to compute a two-dimensional embedding for visualization based on the 30 first principal components (RunUMAP, dims = 1:30). A nearest neighbour graph was constructed (FindNeighbors, dims = 1:30) and unsupervised clustering was performed using the Louvain algorithm (FindClusters) with the resolution parameter set to 0.4. Cell types were assigned based on the expression of known marker genes (Luminal: *Krt8*, *Krt18*, *Cd24a*, *Psca*, *Ly6a*; Basal: *Krt5*, *Trp63*, *Krt14*; Proliferating: *Mki67*, *Birc5*).

The neighbourhood graph correlation-based similarity analysis^67^ was applied to compare cell states in the 2D and 3D scRNA-seq data to a scRNA-seq primary mouse prostate tissue reference. To construct a primary tissue reference, publicly available data from Graham et al.^25^ and Karthaus et al.^23^ was obtained, loaded into R and analysed using Seurat. Low-quality transcriptomes (< 1000 transcripts per cell; <300 & > 5000 genes per cell; <10% mitochondrial count fraction) were removed. Gene counts were log-normalized, and 3000 HVGs were selected. Datasets were integrated by applying the Seurat reciprocal principal component analysis (RPCA) workflow. Nearest neighbour graph construction and computation of a two-dimensional UMAP embedding were based on the 30 first RPCA components. The Louvain algorithm was used for unsupervised clustering at a resolution of 0.7. Cell types were assigned based on marker genes curated from Graham et al. and Karthaus et al.^23,25^. Independent neighbourhood graphs based on transcriptional similarity were constructed for each condition (2D Ctrl, 2D DHT, 3D Ctrl, 3D DHT), as well as the reference using the miloR package (v1.6.0)^68^. For the 2D and 3D organoid data, k = 20 nearest neighbors and 30 PCA dimensions, and for the reference k = 50 nearest neighbors and 50 RPCA dimensions were used. The gene expression profile of each neighbourhood was calculated as the mean expression of the genes within all cells constituting a neighbourhood. For each neighbourhooh, the Pearson correlation of gene expression with all neighborhoods in the reference was computed using scrabbitr R package (v0.1.0)^67^. Pearson correlation calculations were performed on the 3000 most variable genes identified within basal and luminal cells in the reference. The similarity was determined as the maximum correlation between each neighbourhood in a 2D/3D condition and a reference neighborhood.

### *Acp3* (*Ppap*) knockout generation

Mouse prostate organoids were edited by CRISPR-Cas9 system using a protocol described previously^38^. Briefly, two guide RNAs against *Acpp* (sgRNA 1: 5′- TGAGCCGGACAGCAAGCCTC-3′, sgRNA 2: 5′-GAGCCGGACAGCAAGCCTCA-3′) were designed, using Benchling (www.benchling.com), to target the first exon of Acpp. gRNAs were cloned into BbsI linearized pSpCas9(BB)-2A-Puro (PX459) V2.0 plasmid (Addgen #62988). Then the plasmid was transfected into the organoid cells via electroporation using the protocol from Fujii et al.^69^. Briefly, two days before the electroporation, Primocin was removed and Noggin-conditioned media was replaced with recombinant Noggin (100 ng ml−1, 250-38, Peprotech). A day before the electroporation, mouse prostate organoids were treated with 1.25% DMSO. On the day of the electroporation, organoids were shredded mechanically and made into single cells using TrypLE Express Enzyme (12605036, Gibco) as described above. For each plasmid, a total of 1 × 105 cells were resuspended in 100 μl of BTXpress buffer (45-0805, BTX Molecular Delivery Systems) supplemented with RHOKi (Y- 27632; 10 μM, M1817, AbMole) and electroporated with 20 µg of pSPCas9(BB)−2A- Puro V2.0 plasmid (Addgene #62988), containing either gRNA1 or gRNA2, in a nucleofection cuvette (EC-002S, Nepagene) using the NEPA 21 Super Electroporator (Nepagene) with the following settings: Poring pulse (175 V, pulse length—5 ms, pulse interval—50 ms, number of pulses—2, decay rate—10% and positive polarity), Transfer pulse (20 V, pulse length—50 ms, pulse interval—50 ms, number of pulses 5, decay rate—40% and positive/negative polarity).

After electroporation, cells were washed with Opti-MEM (31985070, Gibco) containing RHOKi (Y-27632; 10 μM, M1817, AbMole) and seeded in Matrigel (356231, Corning). The organoid culture medium was supplemented with RHOKi (Y-27632; 10 μM, M1817, AbMole) and DHT (1 nM, A8380, Sigma-Aldrich). Acpp-KO organoids were selected with puromycin (1 µg ml−1, sc-108071B, Santa Cruz Biotechnologies) for 72 h one day after electroporation. Individual organoids were isolated and expanded until molecular characterization. Genomic DNA from potential KO organoid lines was isolated and used to amplify a fraction of *Acpp* containing the sgRNAs (sgRNA 1: 5′- TGAGCCGGACAGCAAGCCTC-3′, sgRNA 2: 5′-GAGCCGGACAGCAAGCCTCA-3′). *Acpp* KO was confirmed by Sanger sequencing and HCR RNA-FISH.

### RNA-fluorescence in situ hybridization (HCR RNA-FISH)

HCR 3.0 Probe Maker v0.3.2 Jupyter Notebook, developed by Özpolat Lab^70^ and publicly available for noncommercial use at https://github.com/rwnull/insitu_probe_generator, was utilised to design the HCR RNA-FISH 3.0 Probes. The sequence of the relevant RNA was used as input by the Python 3 notebook. The mouse genome was blasted with split probes to decrease off-target alignment. Splitprobe couples were eliminated from the pool if the second-highest match was on the same gene and the splitprobe had a match rate greater than 60%. The target sequences were obtained from NCBI (*Acpp*, NM_207668.2).

DNA Probe Sets were reconstituted to a final concentration of 1 µM in TE-Buffer (12090015, Invitrogen). HCR RNA-FISH was performed according to the protocol for mammalian cells from Molecular Instruments, and HCR Amplifier and HCR Buffers sets (Molecular Instruments) were used. Briefly, cells were fixed with 4% PFA for 15 min, and overnight permeabilized with 70% EtOH. After 30 min of prehybridization at 37°C (incubation with hybridization buffer; HCR Buffers set, Molecular Instruments), hybridisation took place overnight with a 1:100 dilution of the probe in hybridization buffer (HCR Buffers set, Molecular Instruments). Samples were washed with heated wash buffer (HCR Buffers set, Molecular Instruments) and three washes with 5xSCCT (20x sodium chloride sodium citrate; 7732-18-5, Invitrogen; + 0.1% Tween in dd H2O). Samples were pre-incubated with amplification buffer (HCR Buffers set, Molecular Instruments) for 30 min. The preferred amplifier (B1 - 488, B2 - 594, B3 – 647; HCR Amplifier set, Molecular instruments) was snap-cooled to 95 degrees and let cool down in the dark for 30 min, then diluted 1:100 in amplification buffer (HCR Buffers set, Molecular Instruments). Amplification was carried out overnight at room temperature. The samples were stained with DAPI (62247; Thermo Fisher, 1:1,000) in 5xSCCT and rinsed in 5x SCCT. Confocal microscopy was used for imaging, as described above.

### Statistical analysis

Unless otherwise indicated, data are presented as mean ± standard deviation (SD), with the exact number of experiments performed indicated in Figure Legends. Statistical analysis was performed using Prism Software (GraphPad). Normal distribution of the data was assessed by the Shapiro-Wilk test. For statistical comparison of datasets from two conditions, two-tailed Student’s t-test was used; for data from three or more conditions/groups, one-way ANOVA with Tukey’s or Dunnett’s post-hoc test was used. Statistical analyses and *p*-values are detailed in Supplementary Table 3.

## DATA AVAILABILITY

scRNA-seq data samples are publicly available in the GEO repository database under the accession number GSE275482 (www.ncbi.nlm.nih.gov/geo/query/acc.cgi?acc=GSE275482). All other relevant data are available from the corresponding authors upon request.

## ACKNOWLEDGEMENTS

This work was supported by the BMBF (FiRe-UPec, 01KI2107 to C.A.), the Deutsche Forschungsgemeinschaft (DFG GRK 2157; 3D Tissue Models for Studying Microbial Infections by Human Pathogens, Project 11, to C.A., Project 12 to A-E.S), and the Single-Cell Center Würzburg (Seed grant to C.A.). A-E.S was supported by the DFG-funded project CRC1583 DECIDE - Decisions in Infectious Diseases; project Z02.

U.D. was supported by the Interdisciplinary Centre for Clinical Research Münster (grant no. Dob2/010/22). We thank Fabian Imdahl (Single cell Center University of Würzburg) for support with scRNA-seq experiments, Natalie Burkard and Nicolas Schlegel (University Hospital Würzburg) for support with ECIS1600R equipment and Stephanie Lamer (University of Würzburg) for Mass Spectrometry analysis. The authors would like to acknowledge critical reading by Vivek Thacker and Mona Alzheimer. Schematics were created with BioRender.com.

## AUTHOR CONTRIBUTIONS

C.A. conceived the study. C.A. wrote the manuscript with the input of all the authors.

M.G., A.J., S.D., S.P. performed most of the experiments in the manuscript. T.B. performed the RNA Fish experiments under the supervision of A-E.S., A.M.L. provided support on the analysis of scRNA-seq data under the supervision of A-E.S.; M.R., C.K. provided technical and material support with the human prostate tissue, U.D. provided technical and material support with the UPEC mutant. All authors contributed to the revision of the manuscript and approved the final version.

## REFERENCES

1. Yang, X. et al. Disease burden and long-term trends of urinary tract infections: A worldwide report. Front. Public Health 10, 888205 (2022).

2. Ulleryd, P. Febrile urinary tract infection in men. Int. J. Antimicrob. Agents 22 **Suppl 2**, 89–93 (2003).

3. Wagenlehner, F. M. E., Weidner, W., Pilatz, A. & Naber, K. G. Urinary tract infections and bacterial prostatitis in men. Curr. Opin. Infect. Dis. 27, 97–101 (2014).

4. Lipsky, B. A. Prostatitis and urinary tract infection in men: what’s new; what’s true? Am. J. Med. 106, 327–334 (1999).

5. Davis, N. G. & Silberman, M. Acute Bacterial Prostatitis. in StatPearls (StatPearls Publishing, 2024).

6. Lupo, F. & Ingersoll, M. A. Is bacterial prostatitis a urinary tract infection? Nat. Rev. Urol. 16, 203–204 (2019).

7. Klein, R. D. & Hultgren, S. J. Urinary tract infections: microbial pathogenesis, host-pathogen interactions and new treatment strategies. Nat. Rev. Microbiol. 18, 211–226 (2020).

8. Lipsky, B. A., Byren, I. & Hoey, C. T. Treatment of bacterial prostatitis. Clin. Infect. Dis. 50, 1641–1652 (2010).

9. Mulvey, M. A. et al. Induction and evasion of host defenses by type 1-piliated uropathogenic Escherichia coli. Science 282, 1494–1497 (1998).

10. Anderson, G. G. et al. Intracellular bacterial biofilm-like pods in urinary tract infections. Science 301, 105–107 (2003).

11. Eto, D. S., Jones, T. A., Sundsbak, J. L. & Mulvey, M. A. Integrin-mediated host cell invasion by type 1-piliated uropathogenic Escherichia coli. PLoS Pathog. 3, e100 (2007).

12. Mysorekar, I. U. & Hultgren, S. J. Mechanisms of uropathogenic Escherichia coli persistence and eradication from the urinary tract. Proc Natl Acad Sci USA 103, 14170–14175 (2006).

13. Shen, M. M. & Abate-Shen, C. Molecular genetics of prostate cancer: new prospects for old challenges. Genes Dev. 24, 1967–2000 (2010).

14. Rudick, C. N. et al. Uropathogenic Escherichia coli induces chronic pelvic pain. Infect. Immun. 79, 628–635 (2011).

15. Quick, M. L. et al. Th1-Th17 cells contribute to the development of uropathogenic Escherichia coli-induced chronic pelvic pain. PLoS ONE 8, e60987 (2013).

16. Boehm, B. J., Colopy, S. A., Jerde, T. J., Loftus, C. J. & Bushman, W. Acute bacterial inflammation of the mouse prostate. Prostate 72, 307–317 (2012).

17. Wong, L., Hutson, P. R. & Bushman, W. Resolution of chronic bacterial-induced prostatic inflammation reverses established fibrosis. Prostate 75, 23– 32 (2015).

18. Simons, B. W. et al. A human prostatic bacterial isolate alters the prostatic microenvironment and accelerates prostate cancer progression. J. Pathol. 235, 478–489 (2015).

19. Elkahwaji, J. E., Zhong, W., Hopkins, W. J. & Bushman, W. Chronic bacterial infection and inflammation incite reactive hyperplasia in a mouse model of chronic prostatitis. Prostate 67, 14–21 (2007).

20. Lupo, F., Rousseau, M., Canton, T. & Ingersoll, M. A. The Immune System Fails to Mount a Protective Response to Gram-Positive or Gram-Negative Bacterial Prostatitis. J. Immunol. 205, 2763–2777 (2020).

21. Khalili, M. et al. Loss of Nkx3.1 expression in bacterial prostatitis: a potential link between inflammation and neoplasia. Am. J. Pathol. 176, 2259–2268 (2010).

22. Love, J. R. & Karthaus, W. R. Next-Generation Modeling of Cancer Using Organoids. Cold Spring Harb. Perspect. Med. 14, (2024).

23. Karthaus, W. R. et al. Regenerative potential of prostate luminal cells revealed by single-cell analysis. Science 368, 497–505 (2020).

24. Drost, J. et al. Organoid culture systems for prostate epithelial and cancer tissue. Nat. Protoc. 11, 347–358 (2016).

25. Graham, M. K. et al. Single-cell atlas of epithelial and stromal cell heterogeneity by lobe and strain in the mouse prostate. Prostate 83, 286–303 (2023).

26. Karthaus, W. R. et al. Identification of multipotent luminal progenitor cells in human prostate organoid cultures. Cell 159, 163–175 (2014).

27. Olson, P. D. et al. Androgen exposure potentiates formation of intratubular communities and renal abscesses by Escherichia coli. Kidney Int. 94, 502– 513 (2018).

28. Mulvey, M. A., Schilling, J. D. & Hultgren, S. J. Establishment of a persistent Escherichia coli reservoir during the acute phase of a bladder infection. Infect. Immun. 69, 4572–4579 (2001).

29. Lane, M. C. & Mobley, H. L. T. Role of P-fimbrial-mediated adherence in pyelonephritis and persistence of uropathogenic Escherichia coli (UPEC) in the mammalian kidney. Kidney Int. 72, 19–25 (2007).

30. Spaulding, C. N. & Hultgren, S. J. Adhesive pili in UTI pathogenesis and drug development. Pathogens 5, (2016).

31. Zhou, G. et al. Uroplakin Ia is the urothelial receptor for uropathogenic Escherichia coli: evidence from in vitro FimH binding. J. Cell Sci. 114, 4095– 4103 (2001).

32. Pak, J., Pu, Y., Zhang, Z. T., Hasty, D. L. & Wu, X. R. Tamm-Horsfall protein binds to type 1 fimbriated Escherichia coli and prevents E. coli from binding to uroplakin Ia and Ib receptors. J. Biol. Chem. 276, 9924–9930 (2001).

33. McLellan, L. K. et al. A host receptor enables type 1 pilus-mediated pathogenesis of Escherichia coli pyelonephritis. PLoS Pathog. 17, e1009314 (2021).

34. Uhlén, M. et al. Tissue-based map of the human proteome. Science 347, 1260419 (2015).

35. Schembri, M. A., Hasman, H. & Klemm, P. Expression and purification of the mannose recognition domain of the FimH adhesin. FEMS Microbiol. Lett. 188, 147–151 (2000).

36. Liu, X. et al. Purification, identification and Cryo-EM structure of prostatic acid phosphatase in human semen. Biochem. Biophys. Res. Commun. 702, 149652 (2024).

37. Martinez, J. J., Mulvey, M. A., Schilling, J. D., Pinkner, J. S. & Hultgren, S. J. Type 1 pilus-mediated bacterial invasion of bladder epithelial cells. EMBO J. 19, 2803–2812 (2000).

38. Aguilar, C. et al. Helicobacter pylori shows tropism to gastric differentiated pit cells dependent on urea chemotaxis. Nat. Commun. 13, 5878 (2022).

39. Blutt, S. E. & Estes, M. K. Organoid models for infectious disease. Annu. Rev. Med. 73, 167–182 (2022).

40. Schutgens, F. & Clevers, H. Human organoids: tools for understanding biology and treating diseases. Annu. Rev. Pathol. 15, 211–234 (2020).

41. Dutta, D. & Clevers, H. Organoid culture systems to study host-pathogen interactions. Curr. Opin. Immunol. 48, 15–22 (2017).

42. Aguilar, C. et al. Organoids as host models for infection biology - a review of methods. Exp. Mol. Med. 53, 1471–1482 (2021).

43. Longhi, C. et al. Features of uropathogenic Escherichia coli able to invade a prostate cell line. New Microbiol. 39, 146–149 (2016).

44. Ho, C.-H. et al. Testosterone regulates the intracellular bacterial community formation of uropathogenic Escherichia coli in prostate cells via STAT3. Int. J. Med. Microbiol. 310, 151450 (2020).

45. Ho, C.-H. et al. Testosterone suppresses uropathogenic Escherichia coli invasion and colonization within prostate cells and inhibits inflammatory responses through JAK/STAT-1 signaling pathway. PLoS ONE 12, e0180244 (2017).

46. Eto, D. S., Sundsbak, J. L. & Mulvey, M. A. Actin-gated intracellular growth and resurgence of uropathogenic Escherichia coli. Cell. Microbiol. 8, 704–717 (2006).

47. Blango, M. G., Ott, E. M., Erman, A., Veranic, P. & Mulvey, M. A. Forced resurgence and targeting of intracellular uropathogenic Escherichia coli reservoirs. PLoS ONE 9, e93327 (2014).

48. Deltourbe, L., Lacerda Mariano, L., Hreha, T. N., Hunstad, D. A. & Ingersoll, M. A. The impact of biological sex on diseases of the urinary tract. Mucosal Immunol. 15, 857–866 (2022).

49. Olson, P. D., Hruska, K. A. & Hunstad, D. A. Androgens enhance male urinary tract infection severity in a new model. J. Am. Soc. Nephrol. 27, 1625– 1634 (2016).

50. 50. Zychlinsky Scharff, A., et al. Sex differences in IL-17 contribute to chronicity in male versus female urinary tract infection. JCI Insight 5, (2019).

51. Hreha, T. N., Collins, C. A., Cole, E. B., Jin, R. J. & Hunstad, D. A. Androgen exposure impairs neutrophil maturation and function within the infected kidney. MBio 15, e0317023 (2024).

52. Weng, Y. et al. Analysis of testosterone and dihydrotestosterone in mouse tissues by liquid chromatography-electrospray ionization-tandem mass spectrometry. Anal. Biochem. 402, 121–128 (2010).

53. Legros, N. et al. PapG subtype-specific binding characteristics of Escherichia coli towards globo-series glycosphingolipids of human kidney and bladder uroepithelial cells. Glycobiology 29, 789–802 (2019).

54. Venegas, M. F. et al. Binding of type 1-piliated Escherichia coli to vaginal mucus. Infect. Immun. 63, 416–422 (1995).

55. Brannon, J. R. et al. Invasion of vaginal epithelial cells by uropathogenic Escherichia coli. Nat. Commun. 11, 2803 (2020).

56. Hiroyama, M. & Takenawa, T. Isolation of a cDNA encoding human lysophosphatidic acid phosphatase that is involved in the regulation of mitochondrial lipid biosynthesis. J. Biol. Chem. 274, 29172–29180 (1999).

57. Tanaka, M. et al. Prostatic acid phosphatase degrades lysophosphatidic acid in seminal plasma. FEBS Lett. 571, 197–204 (2004).

58. Chuang, T.-D. et al. Human prostatic acid phosphatase, an authentic tyrosine phosphatase, dephosphorylates ErbB-2 and regulates prostate cancer cell growth. J. Biol. Chem. 285, 23598–23606 (2010).

59. Zhang, Z., Ostanin, K. & Van Etten, R. L. Covalent modification and site-directed mutagenesis of an active site tryptophan of human prostatic acid phosphatase. Acta Biochim. Pol. 44, 659–672 (1997).

60. Hanley, P. J. Elusive physiological role of prostatic acid phosphatase (PAP): generation of choline for sperm motility via auto-and paracrine cholinergic signaling. Front. Physiol. 14, 1327769 (2023).

61. Quintero, I. B. et al. Prostatic acid phosphatase is not a prostate specific target. Cancer Res. 67, 6549–6554 (2007).

62. Jakob, C. G., Lewinski, K., Kuciel, R., Ostrowski, W. & Lebioda, L. Crystal structure of human prostatic acid phosphatase . Prostate 42, 211–218 (2000).

63. O’Brien, V. P., Hannan, T. J., Nielsen, H. V. & Hultgren, S. J. Drug and vaccine development for the treatment and prevention of urinary tract infections. Microbiol. Spectr. 4, (2016).

64. Langermann, S. et al. Prevention of mucosal Escherichia coli infection by FimH-adhesin-based systemic vaccination. Science 276, 607–611 (1997).

65. Datsenko, K. A. & Wanner, B. L. One-step inactivation of chromosomal genes in *Escherichia coli* K-12 using PCR products. Proc Natl Acad Sci USA 97, 6640–6645 (2000).

66. Bernard, S. C. et al. Pathogenic Neisseria meningitidis utilizes CD147 for vascular colonization. Nat. Med. 20, 725–731 (2014).

67. Ton, M.-L. N. et al. An atlas of rabbit development as a model for single-cell comparative genomics. Nat. Cell Biol. 25, 1061–1072 (2023).

68. Dann, E., Henderson, N. C., Teichmann, S. A., Morgan, M. D. & Marioni, J. C. Differential abundance testing on single-cell data using k-nearest neighbor graphs. Nat. Biotechnol. 40, 245–253 (2022).

69. Fujii, M., Matano, M., Nanki, K. & Sato, T. Efficient genetic engineering of human intestinal organoids using electroporation. Nat. Protoc. 10, 1474–1485 (2015).

70. Kuehn, E. et al. Segment number threshold determines juvenile onset of germline cluster expansion in Platynereis dumerilii. J. Exp. Zool. B Mol. Dev. Evol. 338, 225–240 (2022).

