## Supplementary Material for "Prostate-specific membrane receptor PPAP facilitates *E. coli* invasion of luminal prostate cells via FimH binding"

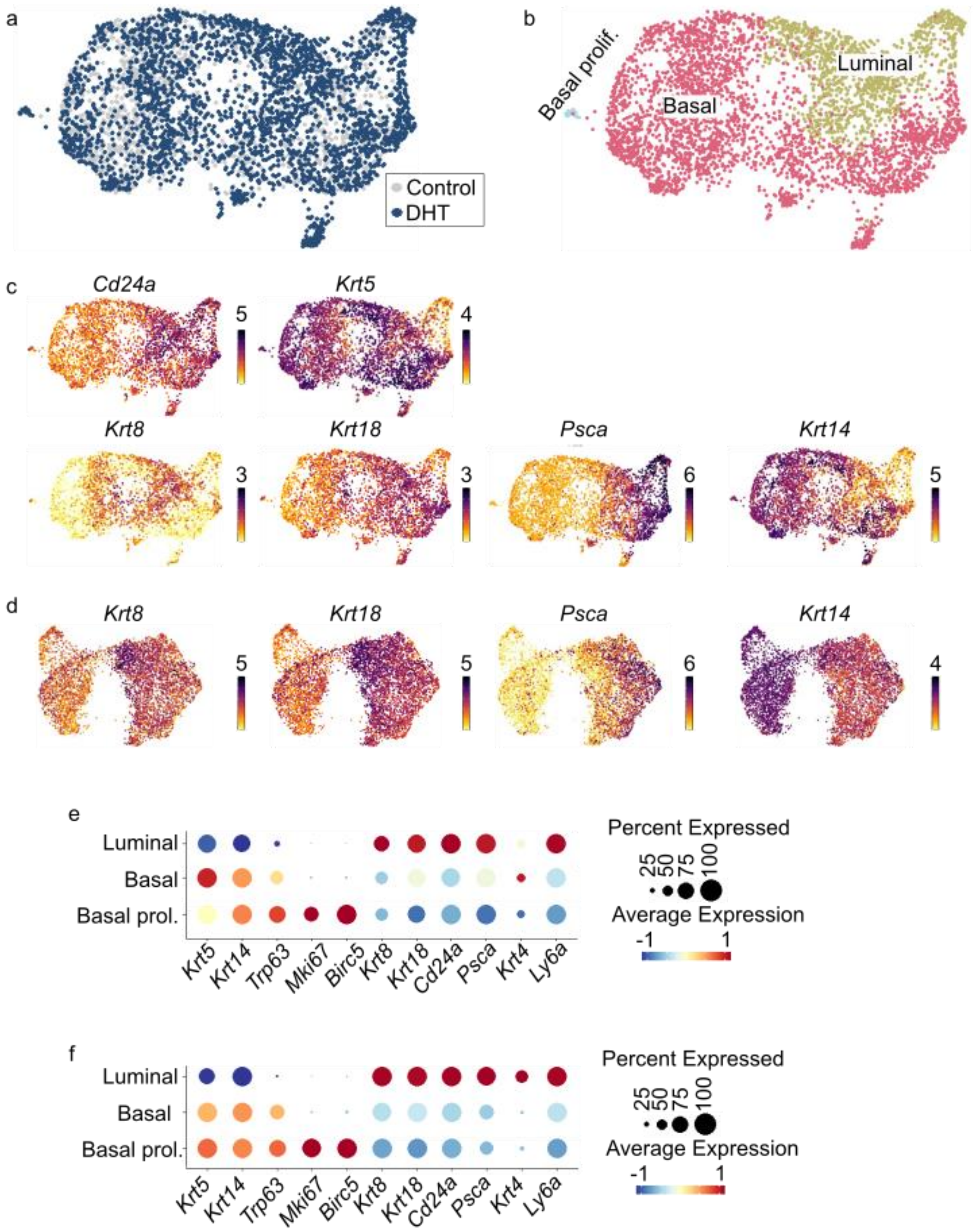


**Extended Data Figure 1. Prostate 3D organoids grown in the absence or presence of DHT (10 nM) contain a high number of basal prostate cells, in contrast to the 2D organoid-based model. a.** Single-cell transcriptomes from the 3D organoids grown in control (grey; 1,471 cells) or DHT (10 nM, dark blue; 2,792 cells) medium were integrated and projected using Uniform Manifold Approximation and Projection (UMAP). **b.** Using the same prostate cell type marker genes as the 2D organoid-based model, cell identities were assigned into three clusters. **c.** Expression of markers specific for luminal prostate cells (*Cd24a, Krt8, Krt18, Psca*) and basal cells (*Krt5, Krt14*) colour-coded and projected on the UMAP displayed in panels (a) and (b). **d.** Expression of markers specific for luminal prostate cells (*Krt8, Krt18, Psca*) and basal cells (*Krt14*) colour-coded and projected on top of the UMAP projection of 2D organoid-based model scRNA-seq data as displayed in Figure 1c-d. **e-f.** Dot plot representation of Z-score of selected marker gene expression on 2D organoid-based model (**e**) and 3D organoids (**f**).

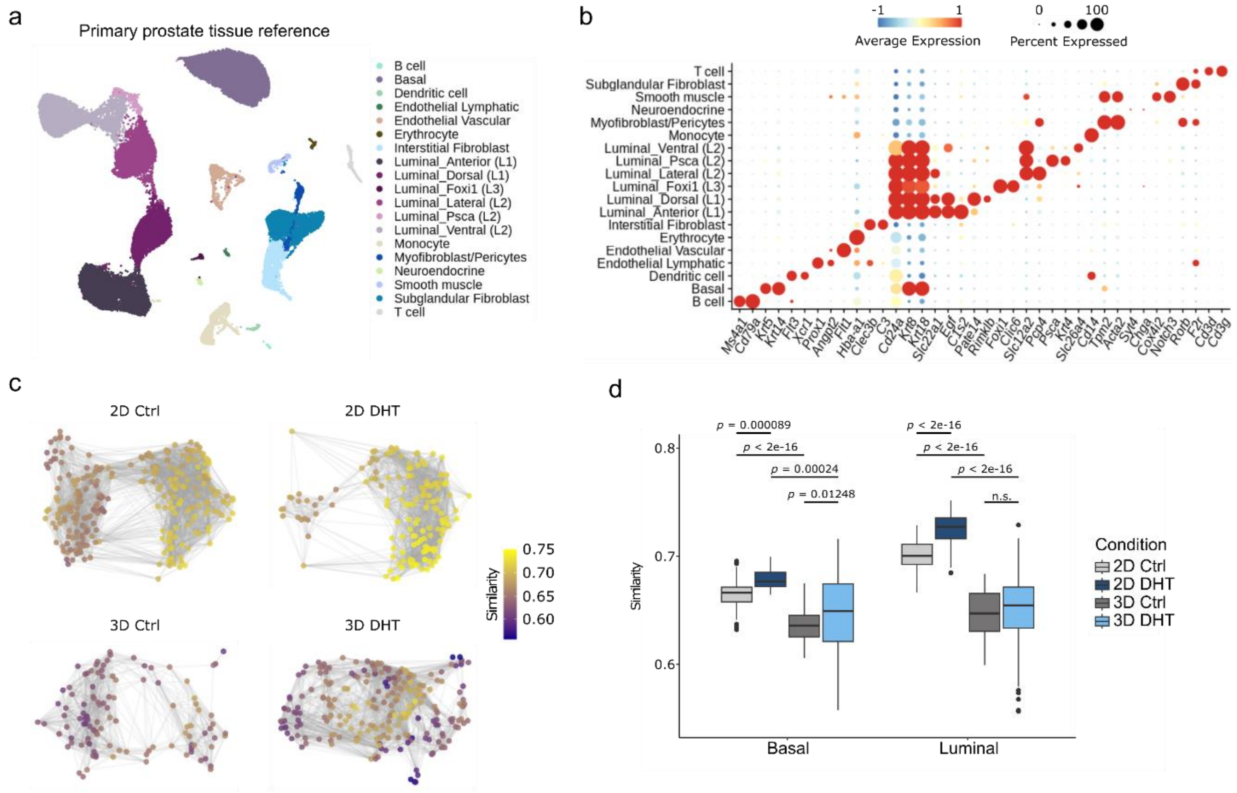


**Extended Data Figure 2. 2D prostate organoid-based model treated with DHT shows the highest similarity to primary prostate tissue on single-cell transcriptomic level. a**. UMAP representation of scRNA-seq transcriptomic data showing the cell types identified in primary prostate tissue scRNA-seq data from two publicly available datasets (Graham et al. 2024, Karthaus et al. 2020). **b**. Dot plot showing scaled mean expression (colour) and percentage of expressing cells (dot size) of selected marker genes used to delineate cell types in the primary tissue reference. **c**. Neighbourhood graphs of the four conditions profiled using scRNA-seq (2D Ctrl, 2D DHT, 3D Ctrl, 3D DHT) coloured by the maximum correlation value across primary prostate reference neighbourhoods. Neighbourhoods are positioned with respect to the UMAP embedding coordinates of the respective index cell. 2D neighbourhoods were constructed from cells across the two replicates in each condition. **d**. Box plot depicting neighbourhood similarities from (**c**) in Basal and Luminal Neighborhoods of 2D/3D. Center-line indicates the median, box limits indicate the upper and lower quantiles, and the whiskers indicate the 1.5x interquartile range.

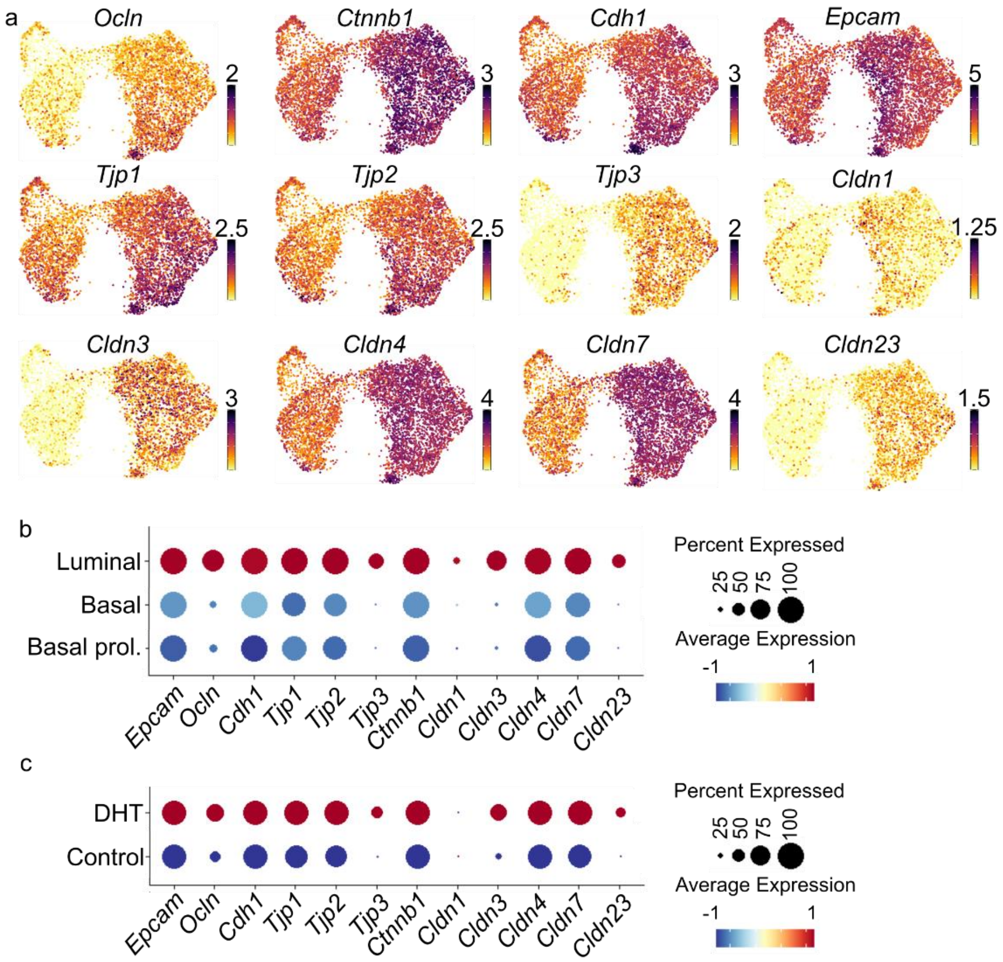


**Extended Data Figure 3. The 2D organoid-based model grown in the presence of DHT (10 nM) express higher levels of barrier function markers. a.** Expression of markers specific for tight junction integrity colour-coded and projected on the UMAP projection of 2D organoid-derived model scRNA-seq data. **b-c.** Dot plot representation of Z-score of selected marker gene expression on 2D organoids based on cell cluster (**b**) and medium treatment (**c**, Control or 10 nM DHT).

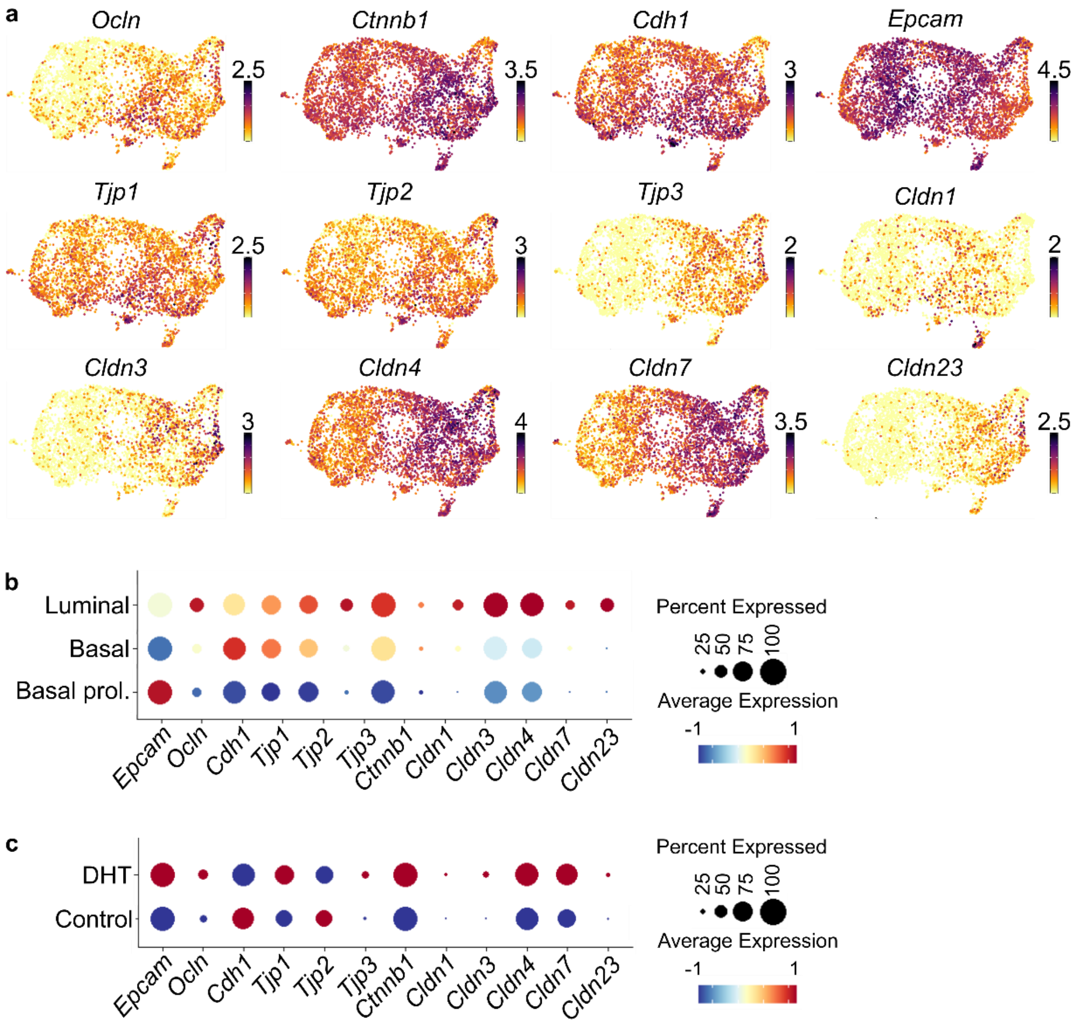


**Extended Data Figure 4. 3D prostate organoids do not express high levels of barrier function markers in the presence of DHT (10 nM). a.** Expression of markers specific for tight junction integrity colour-coded and projected on the UMAP projection of 3D organoids scRNA-seq data. **b-c.** Dot plot representation of Z-score of selected marker gene expression on 3D organoids based on cell cluster (**b**) and medium treatment (**c**, Control or 10 nM DHT).

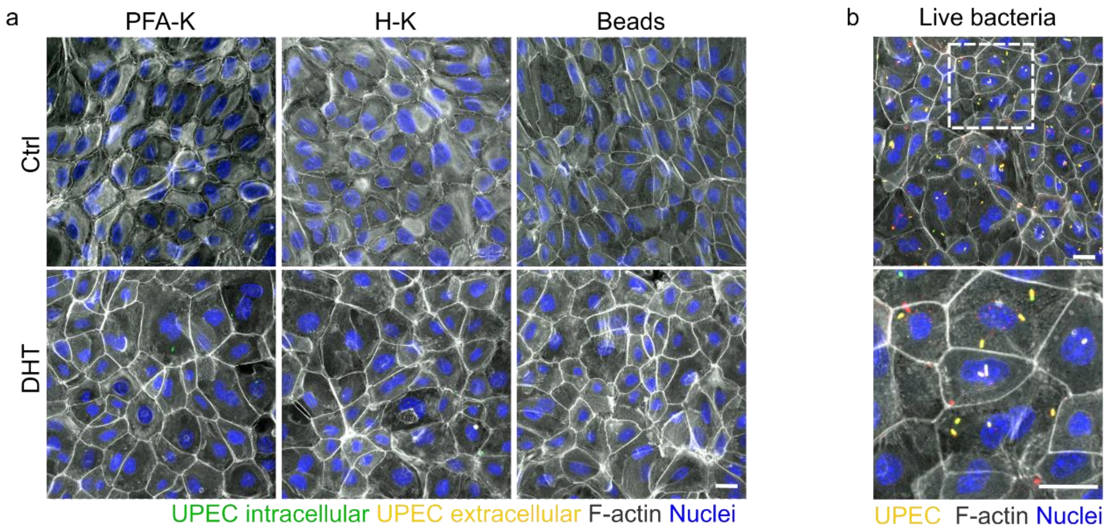


**Extended Data Figure 5. PFA- or Heat-killed UPEC do not invade prostate cells. a.** Representative confocal microscopy images of the organoid-based models incubated with PFA-killed (PFA-K), Heat-killed (H-K) UTI89 or fluorescent beads. F-actin was stained with phalloidin (grey) and nuclei counterstained with Hoechst 33342 (blue). Scale bar 25 µm (*n* = 3). **b.** Experimental control: cells grown in DHT medium were infected with live UPEC expressing GFP and stained with an anti-LPS antibody couple with a secondary Alexa 594 (red). Cells were permeabilised with Triton X-100 (0.5%) prior to staining to confirm that the LPS antibody was working. Nuclei were counterstained using Hoechst 33342 (blue) and F-actin was counterstained using phalloidin (grey). Scale bar 25 µm.

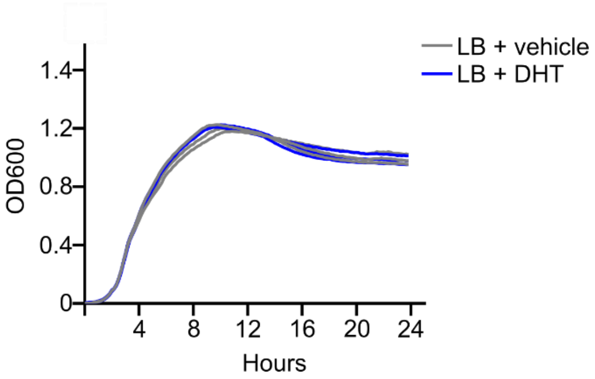


**Extended Data Figure 6. DHT does not affect UPEC growth.** UTI89 growth curve in LB in the presence (blue lines) or absence (grey lines) of 10 nM DHT (*n* = 3).

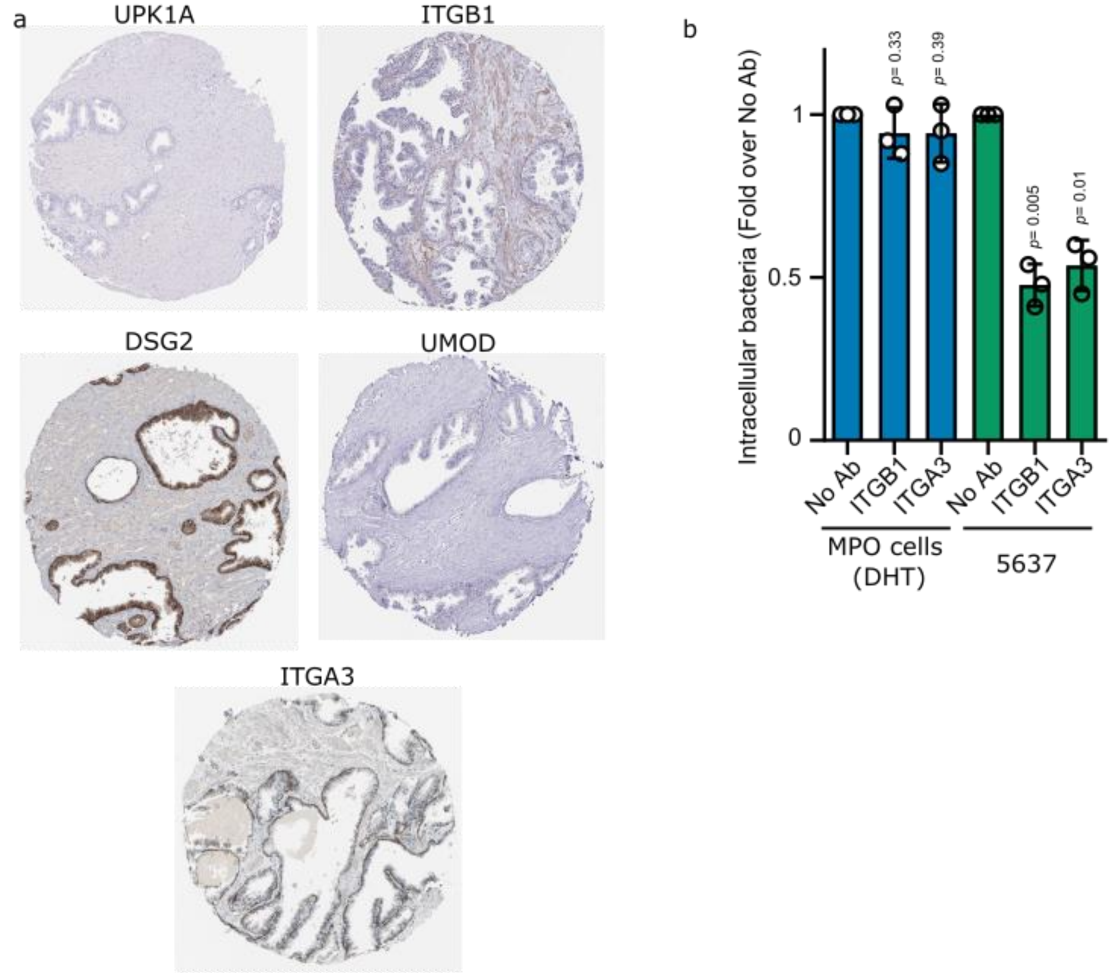


**Extended Data Figure 7. Itgb1 and Itga3 blocking in murine prostate cells does not affect UPEC invasion. a.** Immunohistochemistry analysis of UPK1A, ITGB1, DSG2, UMOD, and ITGA3 protein expression in human prostate tissue (Data from The Human Protein Atlas Database). **b.** Quantification of intracellular bacteria after blocking ITGB1 and ITGA3 with antibodies in the organoid-based model (DHT) or the bladder cell line 5637. No antibody was used as control (No Ab; $n = 3$).

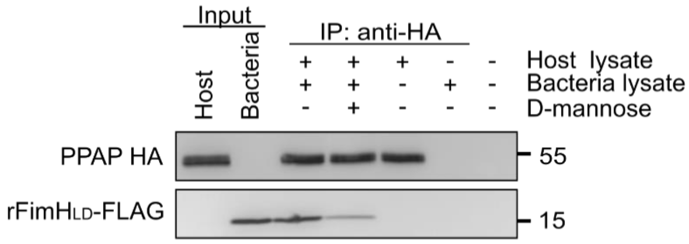


**Extended Data Figure 8. rFimH$_{LD}$-FLAG is co-immunoprecipitated with rPPAP-HA *in vitro*.** Representative image of co-IP of rFimH$_{LD}$-FLAG using the rPPAP-HA as bait (*n* = 3).

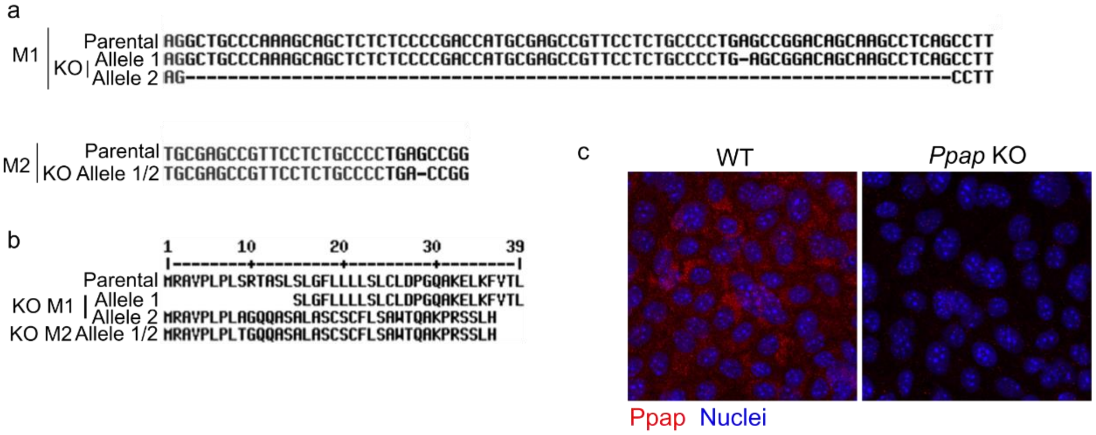


**Extended Data Figure 9. Validation of *Ppap* CRISPR/Cas9 knockout organoid clones**. **a**. Sanger sequencing shows a mutation in *Ppap* sequence for both KO mice (M denotes mouse). **b**. Amino acid sequence shows the sequence for the truncated proteins resulting from the KO. **c**. Representative images of HCR RNA-FISH analysis of *Ppap* on the organoid-based model (grown with DHT) for the WT parental and *Ppap* KO lines ($n$ = 1).
